# Gonadal Hormones Impart Male-Biased Behavioral Vulnerabilities to Immune Activation via Microglial Mitochondrial Function

**DOI:** 10.1101/2022.08.05.502953

**Authors:** Evan A Bordt, Haley A Moya, Young Chan Jo, Caitlin T. Ravichandran, Izabella M. Bankowski, Alexis M. Ceasrine, Christopher J McDougle, William A. Carlezon, Staci D Bilbo

**Affiliations:** Department of Pediatrics, Lurie Center for Autism, Massachusetts General Hospital for Children, Harvard Medical School, Boston, Massachusetts, 02129, USA; Department of Psychology and Neuroscience, Duke University, Durham, North Carolina, 27708, USA; McLean Hospital, Belmont, Massachusetts, 02478, USA; Department of Psychiatry, Harvard Medical School, Boston, Massachusetts, 02115, USA

## Abstract

There is a strong male bias in the prevalence of many neurodevelopmental disorders such as autism spectrum disorder. However, the mechanisms underlying this sex bias remain elusive. Infection during the perinatal period is associated with an increased risk of neurodevelopmental disorder development. Here, we used a mouse model of early-life immune activation that reliably induces deficits in social behaviors only in males. We demonstrate that male-biased alterations in social behavior are dependent upon microglial immune signaling and are coupled to alterations in mitochondrial morphology, gene expression, and function specifically within microglia, the innate immune cells of the brain. Additionally, we show that this behavioral and microglial mitochondrial vulnerability to early-life immune activation is programmed by the male-typical perinatal gonadal hormone surge. These findings demonstrate that social behavior in males over the lifespan are regulated by microglia-specific mechanisms that are shaped by events that occur in early development.

## INTRODUCTION

Neurodevelopmental disorders are a major public health challenge, affecting up to 18% of children worldwide (Lin et al., 2019). There is a strong male bias in the prevalence of early-onset neurodevelopmental disorders such as Autism Spectrum Disorder (ASD) and childhood onset-schizophrenia (SZ) (Goldstein et al., 2013; Halladay et al., 2015; McCarthy and Wright, 2017; Pinares-Garcia et al., 2018; Werling and Geschwind, 2013). For instance, ∼3-4 males are diagnosed with ASD to every female (Baio et al., 2018; Maenner et al., 2020). Despite this strong male bias, the mechanisms that program male vulnerability, or through which females express resilience, remain unknown. Transcriptomic profiling of typically-developing male and female brains demonstrated that genes more highly expressed in male brains were the same genes that were significantly enriched in the brains of individuals diagnosed with ASD (Werling et al., 2016). Interestingly, these same genes were enriched in microglia, the innate immune cells of the central nervous system (Werling et al., 2016). Immune alterations are implicated in the etiology of many neurodevelopmental disorders such as ASD (Inga Jácome et al., 2016; Masi et al., 2015; McDougle et al., 2015). Indeed, maternal infection or fever during the perinatal period is associated with increased risk of offspring ASD diagnoses (Hadjkacem et al., 2016; Hornig et al., 2018; Jiang et al., 2016). Transcriptomic profiling of postmortem brains from individuals diagnosed with ASD identified enrichment of neuroinflammatory genes and dysregulation of genes in microglia (Gandal et al., 2018; Gilman et al., 2011; Quesnel-Vallières et al., 2019; Velmeshev et al., 2019, 2020; Voineagu et al., 2011).

There are striking sex differences in microglial development and function during physiological neurodevelopment (Bordt et al., 2020a; Hanamsagar and Bilbo, 2016; Hanamsagar et al., 2017; VanRyzin et al., 2019a). These developmentally-programmed sex differences are largely organized by the surge of gonadal hormones that occurs only in males during a critical period of perinatal brain development and serves to masculinize the body and brain (McCarthy et al., 2017). Importantly, microglial inhibition prevents hormone-induced masculinization, revealing the importance of microglial function to this male-typical neurodevelopment (Bakker and Baum, 2008; Lenz et al., 2013; Phoenix et al., 1959). Intriguingly, the *in utero* hormonal milieu is hypothesized to play a role in the sex-biased risk profile of ASD (Worsham et al., 2021). Fetal exposure to high levels of the hormone estradiol – the aromatized form of gonadally-produced testosterone (McCarthy, 2009, 2010) - is highly correlated with an increased odds ratio of developing ASD (Baron-Cohen et al., 2020; Bilder et al., 2019). However, other analyses have demonstrated that lower maternal serum levels of unconjugated estriol during the second trimester are also associated with offspring ASD diagnoses (Windham et al., 2016), demonstrating the necessity to understand how gonadal hormones contribute to the sex bias in neurodevelopmental disorder prevalence. Whether gonadal hormone-mediated masculinization of the male brain programs the male bias in neurodevelopmental disorder susceptibility, and the mechanisms through which brain masculinization induces these vulnerabilities, remain unresolved.

Alterations in social behavior are a core symptom of ASD, and microglia play a critical role in the organization of social behaviors in male rodents (Kopec et al., 2018). Microglia are involved in synaptic pruning and refinement, suggesting that microglial dysfunction may contribute to synaptic abnormalities associated with some neurodevelopmental disorders (Paolicelli et al., 2011). Numerous studies suggest an intimate link between microglial inflammatory processes and mitochondrial functions (Bernier et al., 2020a; Nair et al., 2019; York et al., 2020). Importantly, mitochondrial respiratory function is critical for the regulation of microglial homeostatic and inflammatory processes and is impaired following lipopolysaccharide (LPS)-mediated proinflammatory activation (Bernier et al., 2020b; Nair et al., 2019; Orihuela et al., 2016; York et al., 2020). Social behavioral changes may also be regulated by dysfunctional mitochondrial metabolism (Kanellopoulos et al., 2020; Miranda Mendonça et al., 2019; Picard et al., 2018). Mitochondrial metabolism was identified as the key molecular mechanism underlying changes in social behaviors in the CYFIP1 knockout fly model of neurodevelopmental disorders (Kanellopoulos et al., 2020). Postmortem analyses of brain tissues from individuals diagnosed with ASD or SZ revealed decreased expression of mitochondrial electron transport chain (ETC) proteins as well as alterations in proteins regulating mitochondrial morphology (Chauhan et al., 2011; Karry et al., 2004), suggestive of potential deficits in mitochondrial respiratory capacity and function in these brain tissues. Similar to the sex differences in ASD prevalence, sex differences in mitochondrial function in both healthy tissues as well as following perinatal injury have been noted (Demarest et al., 2016b, 2016a; Holody et al., 2021). However, these studies often isolated mitochondria from bulk brain tissue instead of from specific cell types, potentially masking cell-specific sex-specific biology. Whether immune activation during the perinatal period impacts microglial mitochondria, and whether these cell type-specific organellar alterations impact brain function, including social behavior, in a sex-specific manner, is unknown.

Here, we used an established model of early-life immune activation with the bacterial endotoxin LPS (Carlezon et al., 2019; Li et al., 2018; Missig et al., 2018; Patterson, 2011) to investigate whether gonadal hormones present during perinatal brain development induce male vulnerability to immune activation through influencing microglial mitochondrial function. We found that early-life LPS challenge resulted in male-biased deficits in social behavior that were dependent upon microglial immune signaling. This male behavioral vulnerability was coupled to male-biased alterations in microglial mitochondrial morphology, gene expression, and respiratory function. We demonstrated that this vulnerability can be induced by masculinizing female mouse pups at birth with the gonadally-derived hormone estradiol prior to immune activation. These data uncover a fundamental mechanism through which physiological developmental programs may regulate sex- and cell-specific neurological susceptibilities.

## RESULTS

### Early-life immune challenge induces male-specific alterations in social behavior

To assess whether early-life LPS challenge resulted in sex-specific changes in social behavior, we injected female and male mice with LPS at postnatal day 9 (PN9), a time point previously established to induce long-term behavioral changes in male mice (Carlezon et al., 2019). LPS injection decreased body weights at PN10 in both sexes compared to saline-injected controls, but group differences were no longer detectable by PN30 (Supplemental Fig. 1A). We assessed social preference starting at PN30 using a modified Crawley’s 3-chambered arena task (Smith et al., 2015) in which mice were allowed to investigate a novel age- and sex-matched conspecific in one of the side chambers, or a novel object (i.e. small rubber duck) in the opposite chamber (Fig. 1A). Consistent with the juvenile drive to seek out novel social experiences, saline-treated female and male mice both preferentially investigated a novel social stimulus compared to a novel object (Fig. 1B-C, Supplemental Fig. 1C-E). LPS treatment did not alter the time that females spent investigating either the social stimulus or the novel object. However, male mice treated on PN9 with LPS showed decreased social preference at PN30, with decreased social investigation and increased object investigation, and no longer preferred to investigate a novel animal over a novel object (Fig. 1B-C, Supplemental Fig. 1C-E). To measure the preference of a mouse to investigate a novel mouse or a familiar mouse (novelty preference), we then performed a similar 3-chambered task in which the subject mouse is placed in a 3-chambered arena and given the option to investigate either a new novel age- and sex-matched conspecific or a familiar cage mate animal (Fig. 1D). Consistent with alterations in sociability, saline-treated mice preferred to investigate the novel mouse over their cage-mate sibling, and LPS treatment did not alter social novelty preference in female mice. Treatment with LPS at PN9 decreased novel investigation, increased investigation of the familiar animal, and decreased social novelty preference in ∼PN30 males (Fig. 1E-F, Supplemental Fig 1F-H). While early-life immune challenge induced male-biased alterations in several social behavior tasks, no changes were observed in the elevated zero maze (Fig. 1G), light dark box test (Fig. 1H), or in marble burying (Supplemental Fig. 1B), suggesting that changes in anxiety-like behaviors or repetitive behaviors do not explain deficits in social behaviors.

**Figure 1.**
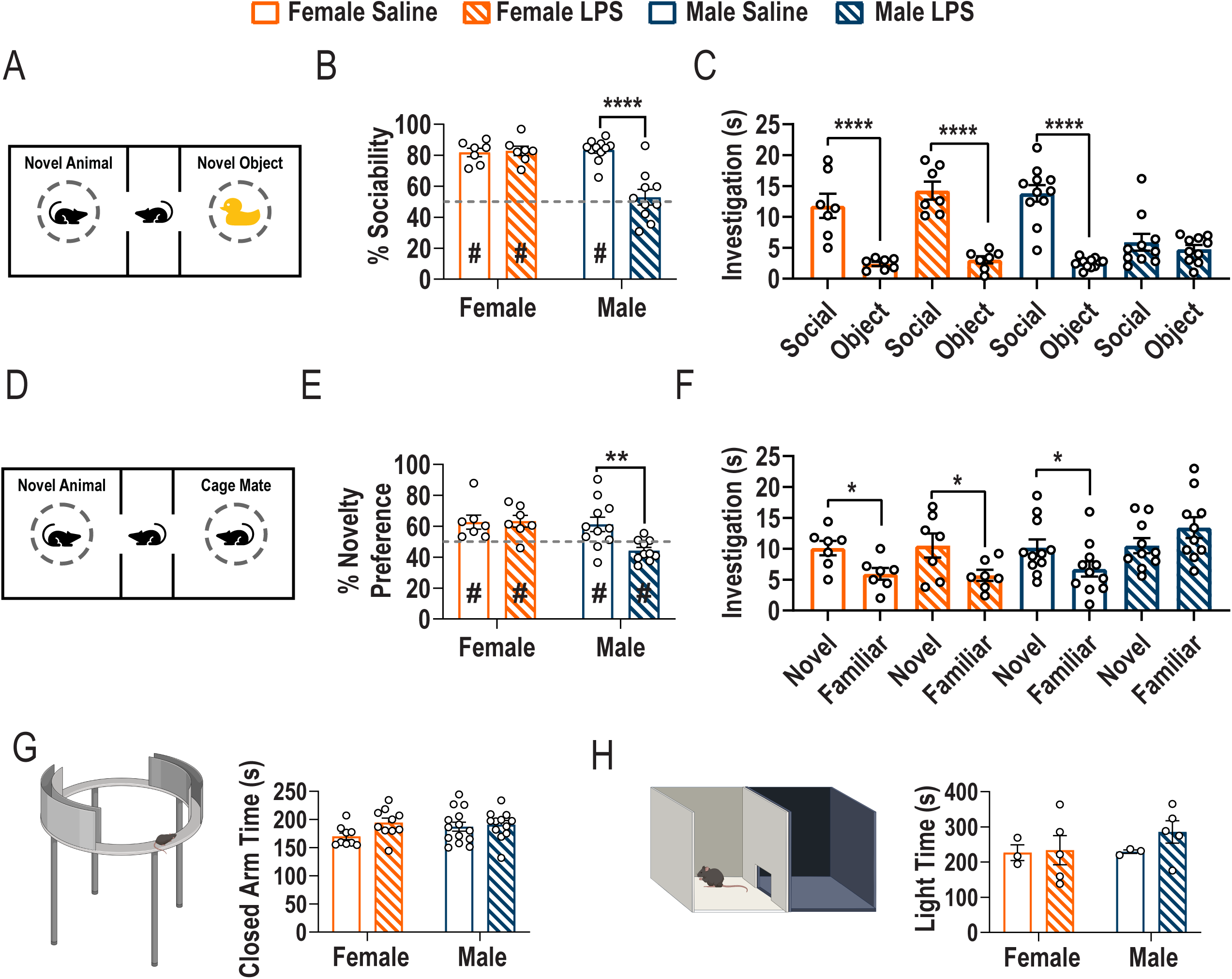
Perinatal immune challenge induces deficits in social behavior in male but not female mice. **A**. PN30 female (orange) and male (blue) mice challenged with saline (Sal: open bars) or LPS (striped bars) at PN9 were placed in a 3-chambered arena and given the choice to interact with a novel conspecific mouse or an inanimate object for 5 min. **B**. Time investigating the novel animal compared to novel object (% sociability) was assessed. Differences across groups were assessed by 2-way ANOVA followed by Bonferonni’s post-hoc analyses. **** p < 0.0001. # single sample t-test against 50% (chance), p < 0.05. Bar graphs display mean ± standard error of the mean. **C**. Social investigation time and object investigation time was compared within saline-treated and LPS-treated female and male mice. Differences between social and object investigation time were assessed by paired t-tests. **** p < 0.0001. Bar graphs display mean ± standard error of the mean. **D**. PN30 female (orange) and male (blue) mice challenged with saline (Sal: open bars) or LPS (striped bars) at PN9 were placed in a 3-chambered arena and given the choice to interact with a sex- and age-matched novel conspecific or a cage mate for 10 min. **E**. Time spent investigating the novel animal compared to cage mate (% novel investigation) was assessed. Differences across groups were assessed by 2-way ANOVA followed by Bonferonni’s post-hoc analyses. ** p < 0.01. # single sample t-test against 50% (chance), p < 0.05. Bar graphs display mean ± standard error of the mean. **F**. Novel investigation time and familiar investigation time was compared within saline-treated and LPS-treated female and male mice. Differences between novel and familiar investigation time were assessed by paired t-tests. **** p < 0.0001. Bar graphs display mean ± standard error of the mean. **G**. ∼PN30 mice were placed in an elevated zero maze for 5 min and time spent in closed arms was assessed. Differences across groups were assessed by 2-way ANOVA. Bar graphs display mean ± standard error of the mean. Figure was created using Biorender. **H**. ∼PN30 mice were placed in a light dark box for 10 min and time spent in the light zone was assessed. Differences across groups were assessed by 2-way ANOVA. Bar graphs display mean ± standard error of the mean. Figure was created using Biorender.

### Social behavior differences are not caused by sex-biased monocyte infiltration

Given the known ability of LPS injection to induce trafficking of peripheral leukocytes into the brain (Cazareth et al., 2014), we next tested the hypothesis that sex differences in social behavior in response to LPS challenge are due to elevated infiltration of peripheral monocytes into the brain in males. We collected prefrontal cortex (PFC) tissue, based on the relevance of this brain region to social behavior and its disruption in disorders such as ASD, from female and male mice at 24 and at 72 hr following saline or LPS challenge and performed flow cytometry to determine the extent of brain infiltration. We defined microglia in the PFC as CD11b^+^CD45^lo^ cells, and infiltrating monocytes as CD11b^+^CD45^hi^ cells (Philpott et al., 2022)(Supplemental Fig. 2A). CD11b^+^CD45^hi^ cells express the chemokine receptor CCR2, whereas CD11b^+^CD45^lo^ cells do not express CCR2 (Supplemental Fig. 2B). CD11b^+^CD45^lo^ microglia also expressed the fractalkine receptor Cx3cr1, whereas CD11b^+^CD45^hi^CCR2^+^ infiltrating monocytes did not express Cx3cr1 (Fig. 2C). We observed no sex differences in Cx3cr1 expression within either CD11b^+^CD45^lo^ microglia or CD11b^+^CD45^hi^ infiltrating cells (Supplemental Fig. 2C). Consistent with previous reports, we observed that LPS challenge induced infiltration of peripheral monocytes into the brain 72 hr following injection, although we found negligible infiltration 24 hr following LPS injection and no change in microglial number at either time point (Fig. 2A-B, Supplemental Fig. 2D). In sum, there were no sex differences in brain infiltration of CD11b^+^CD45^hi^ cells (Fig. 2B), suggesting that the sex differences observed in social behavior following LPS injection are not caused by male-biased infiltration of peripheral monocytes into the brain.

**Figure 2.**
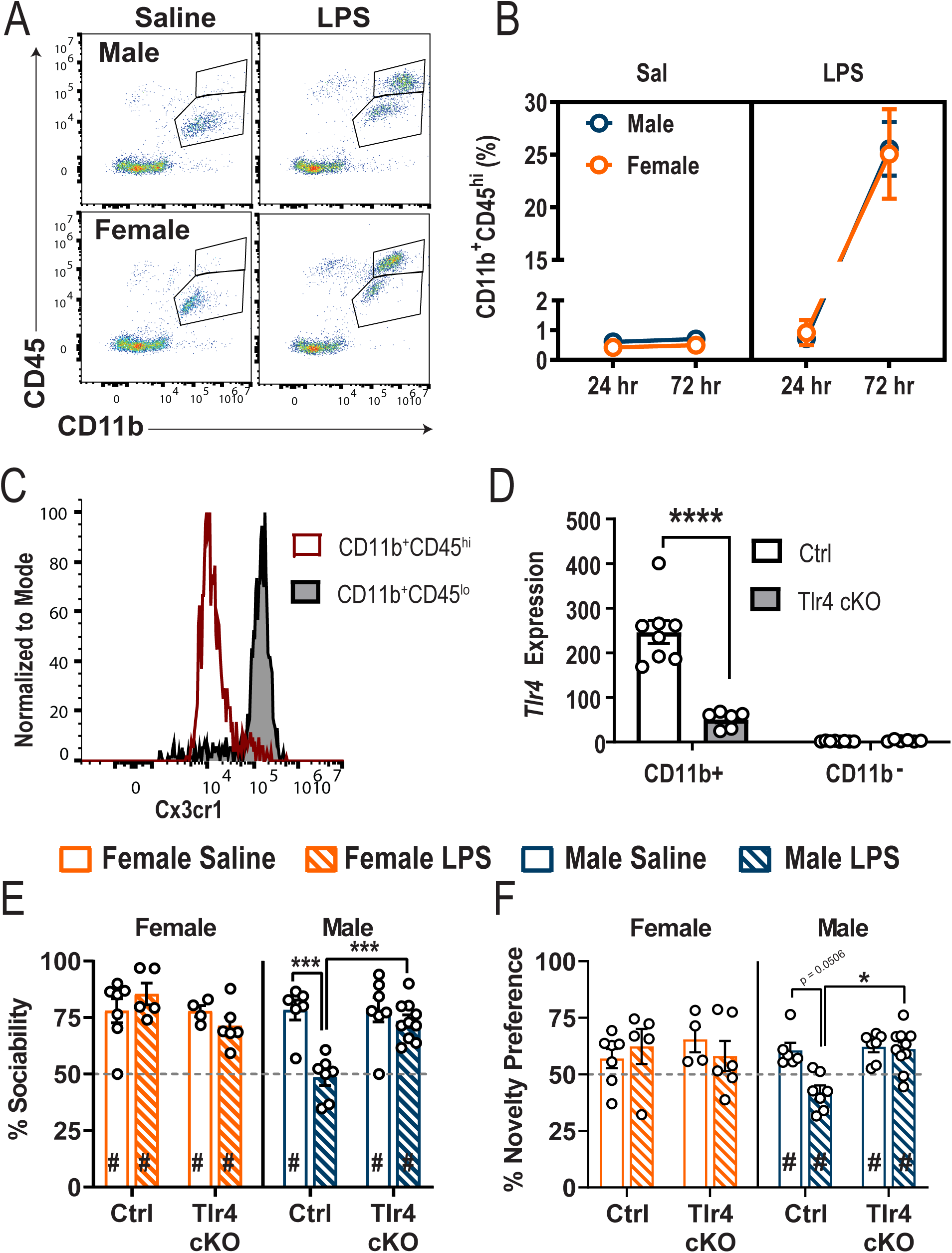
Male-biased social behavior deficits are dependent upon microglial Tlr4. **A**. Anterior cingulate cortex (ACC) from female and male mice injected on PN9 with saline or LPS and collected either 24 hr or 72 hr post-injection. ACC tissue was dissociated, brought to a single cell suspension, and cleaned with Miltenyi Debris Removal Solution prior to flow cytometric analysis. Tissue was analyzed for expression of CD11b/CD45. Cells were gated on forward scatter (FSC)/side scatter (SSC), singlets, live, CD11b, CD45, CCR2, and Cx3cr1. **B**. Brain cells from PN10 and PN12 male and female mice injected on PN9 with saline or LPS were analyzed for CD11b positivity and CD45 hi/lo status. The percentage of cells assessed to be CD11b^+^CD45^hi^ is presented. Graphs display mean ± standard error of the mean. **C**. Cx3cr1 expression was assessed within CD11b^+^CD45^lo^ and CD11b^+^CD45^hi^ cell populations. **D**. Microglia (CD11b+ cells) and non-microglia (CD11b^-^ cells) were isolated from the prefrontal cortex of Tlr4flox/flox (Ctrl) and Cx3cr1-CrebT:Tlr4flox/flox (Tlr4 cKO) mice. qPCR for *Tlr4* relative to *18S* was performed to confirm knockdown of *Tlr4* expression in Cx3cr1+ cells. Differences across groups were assessed by 2-way ANOVA followed by Bonferonni’s post-hoc analyses. Bar graphs display mean ± standard error of the mean. **** p < 0.0001. **E**. PN30 female (orange) and male (blue) Ctrl and Tlr4 cKO mice challenged with saline (Sal: open bars) or LPS (striped bars) at PN9 were placed in a 3-chambered arena and given the choice to interact with a novel conspecific mouse or an inanimate object for 5 min. Time investigating the novel animal compared to novel object (% sociability) was assessed. Differences across groups were assessed by 3-way ANOVA followed by Bonferonni’s post-hoc analyses. *** p < 0.001, **** p < 0.0001. # single sample t-test against 50% (chance), p < 0.05. Bar graphs display mean ± standard error of the mean. **F**. PN30 female (orange) and male (blue) Ctrl and Tlr4 cKO mice challenged with saline (Sal: open bars) or LPS (striped bars) at PN9 were placed in a 3-chambered arena and given the choice to interact with a sex- and age-matched novel conspecific or a cage mate for 10 min. Time spent investigating the novel animal compared to cage mate (% novel investigation) was assessed. Differences across groups were assessed by 3-way ANOVA followed by Bonferonni’s post-hoc analyses. * p < 0.05. # single sample t-test against 50% (chance), p < 0.05. Bar graphs display mean ± standard error of the mean.

### Microglial Tlr4 signaling is required for male-biased social behavioral alterations in response to early-life immune challenge

Considering the lack of sex difference in peripheral monocyte infiltration into the brain following PN9 LPS challenge, we next tested whether early-life immune challenge led to male-specific alterations in social behavior through microglial immune signaling. To assess this question, we bred mice in which the gene for toll-like receptor 4 (Tlr4), a pattern recognition receptor that initiates inflammatory cascades in response to LPS, was flanked by loxP excision sites (floxed) with BAC transgenic Cx3cr1-CreBT mice in order to ablate Tlr4 signaling within microglia throughout the brain (Ceasrine et al., 2022) (Fig. 2D). We then administered a PN9 LPS challenge and assessed social behaviors in these mice lacking microglial Tlr4. Importantly, ablation of Tlr4 signaling within microglia (Tlr4 conditional knockout: Tlr4 cKO) had no effect on sociability or social novelty preference in saline-treated female or male mice nor LPS-treated female mice. However, microglial Tlr4 ablation prevented the LPS-induced deficit in sociability and social novelty preference in male mice (Fig. 2E-F, Supplemental Fig. 2E-J), indicating that microglial immune signaling is necessary for the male-specific deficits in social behaviors induced by early-life immune challenge.

### Perinatal sex hormones organize male behavioral vulnerability to early-life immune challenge

Upon establishing that microglial immune signaling is necessary for the male-biased alterations in social behavior in response to PN9 LPS, we next set out to determine the mechanisms by which this early-life sex difference in microglial susceptibility is programmed. One well-characterized sex-specific developmental program that may underlie this male bias is a surge of gonadal hormones that occurs only in male mice during a perinatal critical period of brain development and organization. This gonadal hormone surge in which the fetal testis produces androgens in the latter third of gestation in rodents (McCarthy et al., 2017; Zuloaga et al., 2008) masculinizes the male body and brain in a microglia-dependent manner (Bakker and Baum, 2008; Lenz et al., 2013; Phoenix et al., 1959). During early postnatal life, the female rodent brain is sensitive to gonadal hormone exposure, and can be diverted towards a male-like state (“masculinized”) through exogenous administration of the gonadal hormone testosterone or its aromatized form estradiol (McCarthy et al., 2017, 2018; VanRyzin et al., 2019b). We predicted that this hormone-driven organization of the male brain imparts vulnerabilities to subsequent immune challenges. To test this hypothesis, we masculinized the female brain by injecting female mouse pups with estradiol on PN1 and PN2 (Fig. 3A). This allows us to determine whether brain organizational changes induced by the perinatal gonadal hormone surge impart male vulnerability to immune challenge independent of the chromosomal milieu. Female mice masculinized with estradiol at birth (F+E_2_) and then injected on PN9 with saline showed typical social drive, preferring the novel social stimulus compared to the novel object, similar to saline-treated females and males. Consistent with our previous data, PN9 LPS induced deficits in sociability and social novelty preference in male but not female mice (Fig. 3B-C, Supplemental Fig. 3D-I). Strikingly, female mice masculinized at birth with estradiol and then challenged at PN9 with LPS showed similar deficits in sociability and social novelty preference as LPS-treated males (Fig. 3B-C, Supplemental Fig. 3D-I). We also assessed an additional social behavior using a juvenile social exploration assay in which PN15 pups were presented with home-cage bedding or novel (age- and treatment-matched) bedding, a test in which neurotypically developing mice prefer to investigate novel bedding (Bilbo et al., 2018)(Supplemental Fig. 3A). We found that PN9 LPS interrupted the exploratory drive, and that masculinizing females at birth introduced a similar susceptibility (Supplemental Fig. 3B-C). These data reveal unique immune challenge-induced alterations in social behaviors that are organized by the perinatal gonadal hormone surge typical of male neurodevelopment.

**Figure 3.**
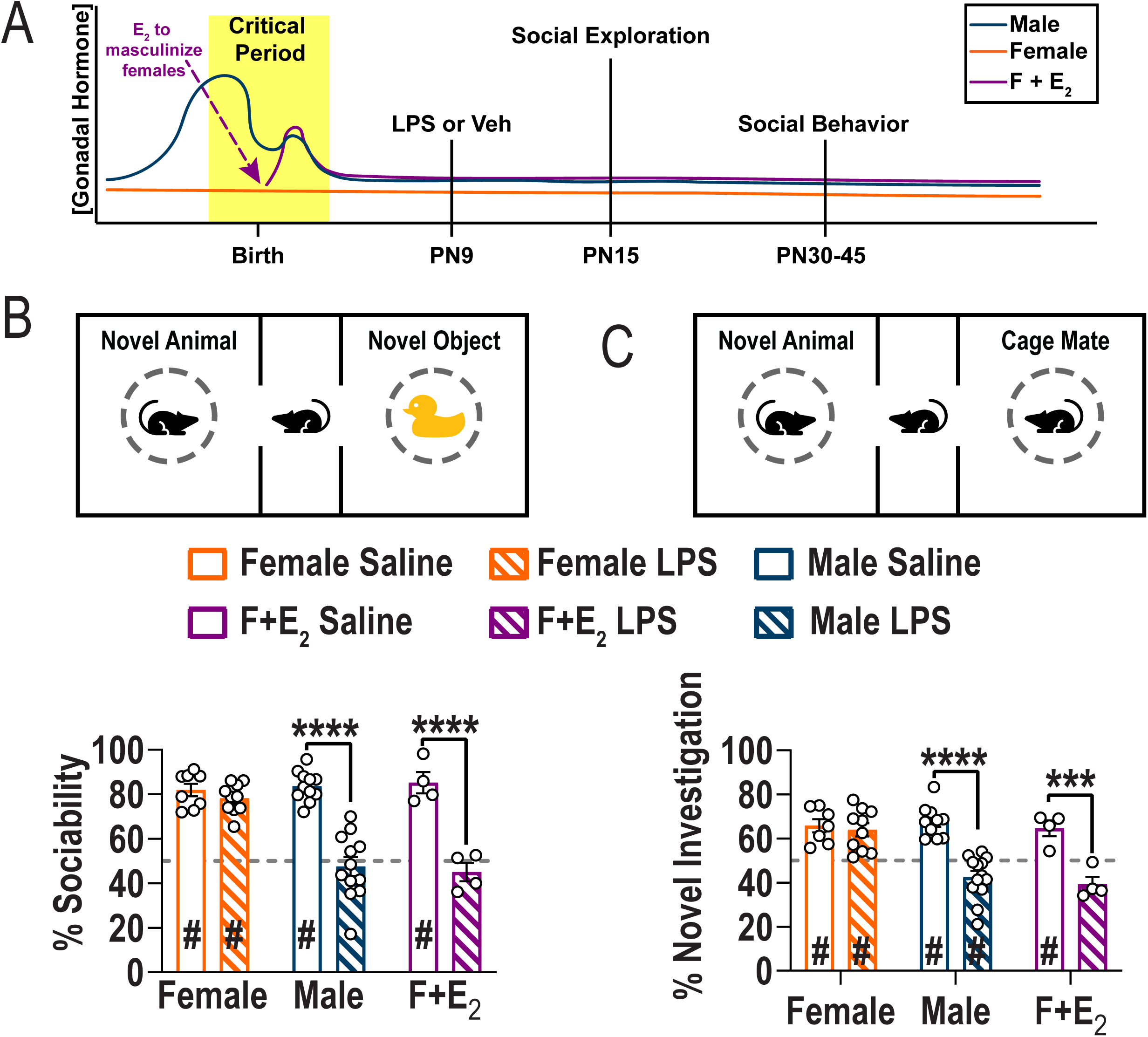
Perinatal sex hormones induce social behavior vulnerability to early life immune challenge. **A**. Female mice are masculinized on PN1-2 by injection with estradiol benzoate (E_2_). Male, female, and masculinized female (F+E_2_) mice are injected on PN9 with saline or LPS. **B**. PN30 female (orange), male (blue), and masculinized female (F+E_2_: purple) mice challenged with saline (Sal: open bars) or LPS (striped bars) at PN9 were placed in a 3-chambered arena and given the choice to interact with a novel conspecific mouse or an inanimate object for 5 min. Time investigating the novel animal compared to novel object (% sociability) was assessed. Differences across groups were assessed by 2-way ANOVA followed by Bonferonni’s post-hoc analyses. **** p < 0.0001. # single sample t-test against 50% (chance), p < 0.05. Bar graphs display mean ± standard error of the mean. **C**. PN30 female (orange), male (blue), and masculinized female (F+E_2_: purple) mice challenged with saline (Sal: open bars) or LPS (striped bars) at PN9 were placed in a 3-chambered arena and given the choice to interact with a sex- and age-matched novel conspecific or a cage mate for 10 min. Time spent investigating the novel animal compared to cage mate (% novel investigation) was assessed. Differences across groups were assessed by 2-way ANOVA followed by Bonferonni’s post-hoc analyses. *** p < 0.001. **** p < 0.0001. # single sample t-test against 50% (chance), p < 0.05. Bar graphs display mean ± standard error of the mean.

### Early-life immune challenge induces long-term male-biased alterations in expression of mitochondrial electron transport chain genes

Our data demonstrate that early-life immune challenge causes male-specific, microglial Tlr4-dependent social behavior alterations that are dependent on the perinatal gonadal hormone surge. We therefore determined what microglial functions were altered by the perinatal gonadal hormone surge that imparts male behavioral vulnerability to immune challenge. Given the known neuroimmune and mitochondrial abnormalities present in early-onset neurodevelopmental disorders, and the recently appreciated link between microglial mitochondrial function and social behavior revealed in male mice, we analyzed our previously published (Hanamsagar et al., 2017) bulk RNA sequencing of isolated microglia treated acutely with LPS for mitochondrial electron transport chain gene expression. In this experiment, PN60 female and male mice were injected with LPS and 2 hr later microglial isolations were performed followed by RNA sequencing (Supplemental Fig. 4A). Sequencing revealed robust male-specific downregulation of mitochondrial electron transport chain gene expression within CD11b+ cells (microglia)(Supplemental Fig. 4B). We next assessed whether this robust sex-specific transcriptomic alteration was simply a kinetic difference in microglial mitochondrial gene expression between the sexes, or whether PN9 LPS treatment would induce lasting male-specific alterations in mitochondrial gene expression on a timescale consistent with the changes in social behavior (i.e. at PN30). Microglia were isolated from dissected prefrontal cortex of PN30 female, male, and masculinized female mice that had been injected on PN9 with saline or LPS and mitochondrial electron transport chain gene expression was assessed by PCR Array (Fig. 4A). Similar to what we observed following acute immune challenge, we observed no decrease in the expression of mitochondrial electron transport chain genes in females treated with LPS compared to saline-treated females (Fig. 4B). PN9 LPS challenge resulted in dramatic decreases in mitochondrial electron transport chain gene expression in microglia isolated from PN30 males compared to saline-treated males (Fig. 4B). Microglia from LPS-treated masculinized females also significantly downregulated mitochondrial gene expression compared to saline-treated mice (Fig. 4B). Interestingly, the majority of mitochondrial electron transport chain genes that remained downregulated by PN30 encode proteins in Complex I, as 94% of Complex I genes were significantly downregulated in LPS-treated male mice, and 91% of these same genes were downregulated in LPS-treated masculinized female microglia (Fig. 4B). Importantly, this mitochondrial vulnerability to perinatal immune challenge appears to be specific to microglia, as there was no significant LPS-induced change in gene expression in CD11b-cells (Supplemental Fig. 4C-D).

**Figure 4.**
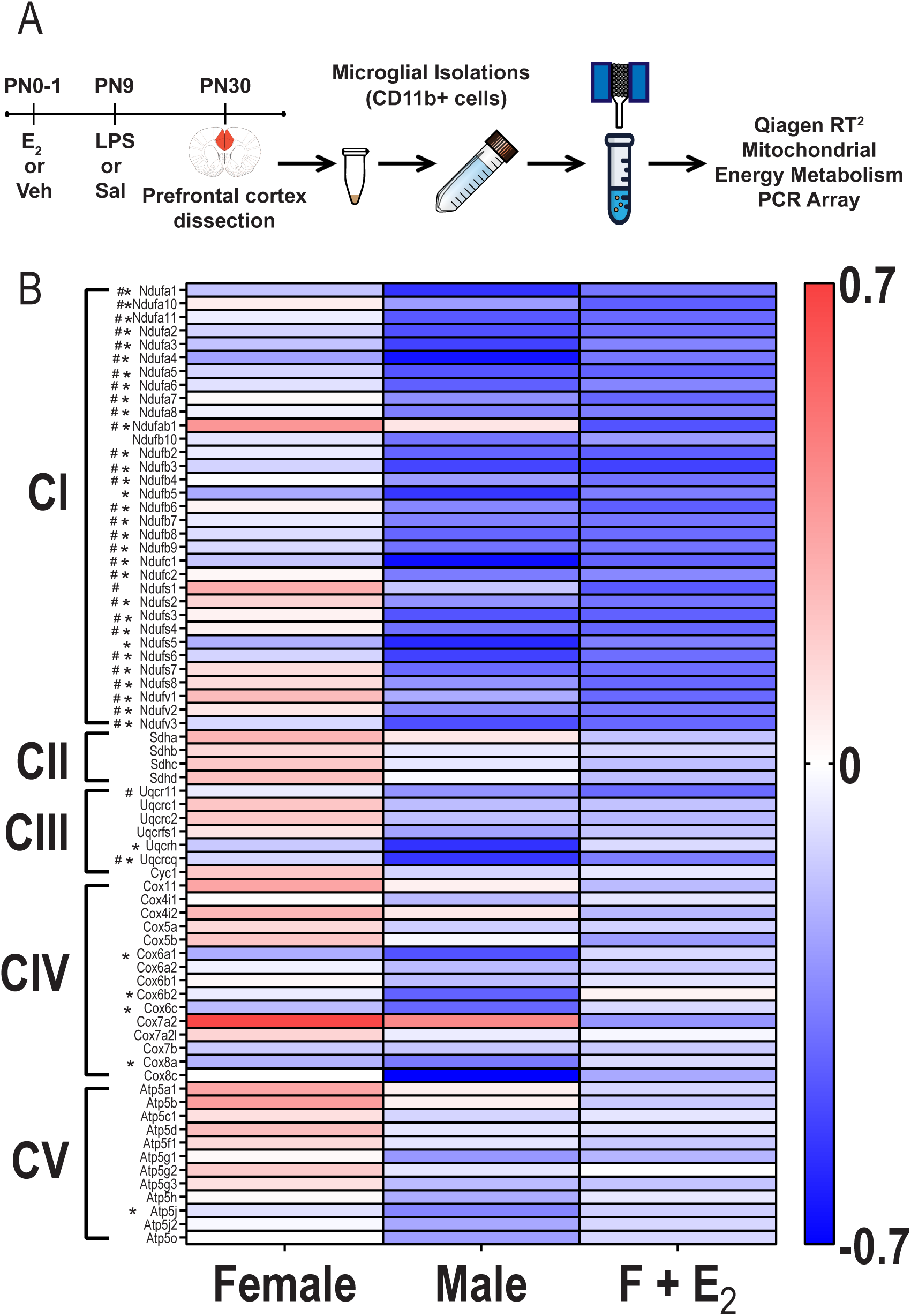
Perinatal immune challenge leads to lasting changes in expression of microglial mitochondrial electron transport chain genes in male and masculinized female - but not in female - mice. **A**. Experimental timeline. Female, male, or females masculinized on PN1-2 with estradiol benzoate were injected on PN9 with LPS or saline vehicle. At PN30, anterior cingulate cortex was dissected, and microglia were isolated by CD11b bead method. Extracted RNA from CD11b+ cells (microglia) was then assessed for mitochondrial electron transport chain (ETC) gene expression using Qiagen RT2 Mitochondrial Energy Metabolism PCR Array. **B**. Log of mRNA expression of ETC subunits (Complexes I-V: CI-CV) from CD11b+ cells isolated from ACC of PN30 mice injected with LPS on PN9 relative to that gene’s expression from saline-treated mice (red = upregulation, white = no change, blue = downregulation). F+E_2_ = masculinized female. Differences across groups were assessed by 2-way ANOVA followed by Bonferonni’s post-hoc analyses. * p < 0.05 for male LPS compared to male saline. # p < 0.05 for F+E_2_ LPS compared to F+E2 saline. There were no LPS-induced differences in female mice.

### Perinatal immune challenge induces lasting male-biased alterations in microglial mitochondrial morphology and function

Given the striking downregulation of mitochondrial electron transport chain genes in microglia from male and masculinized female mice in response to early-life immune challenge, we next assessed whether this early-life immune challenge would result in further sex-biased alterations in microglial mitochondrial function. Changes in mitochondrial architecture through the opposing processes of fission and fusion impact mitochondrial bioenergetic functions (Huertas et al., 2019; Liesa and Shirihai, 2013; Liesa et al., 2009; Wai and Langer, 2016). We assessed mitochondrial morphology within microglia in the anterior cingulate cortex (ACC), a region of the frontal cortex critical for social behaviors. To do so, we stained for the outer mitochondrial membrane protein Tomm20 and the microglial marker ionized calcium binding adaptor molecule 1 (Iba1) from PN30 mice challenged on PN9 with saline or LPS (Fig. 5A). We used volumetric reconstruction software to mask mitochondrial signal within and outside of microglia, and assessed morphology of these reconstructed mitochondria (Chandra et al., 2017). In females, microglial mitochondrial length did not differ regardless of early life treatment. However, PN9 LPS significantly decreased mitochondrial length within male microglia compared to saline-treated males (Fig. 5B-C, Supplemental Fig. Fig. 5A). Microglial mitochondria within the PN30 ACC were also shorter in masculinized females challenged with LPS at PN9 (Fig. 5B-C, Supplemental Fig. 5A). Similar results were found for mitochondrial volume measures, as PN9 LPS treatment led to smaller microglial mitochondria within males and masculinized females, but not in females (Fig. 5D-E, Supplemental Fig. 5B).

**Figure 5.**
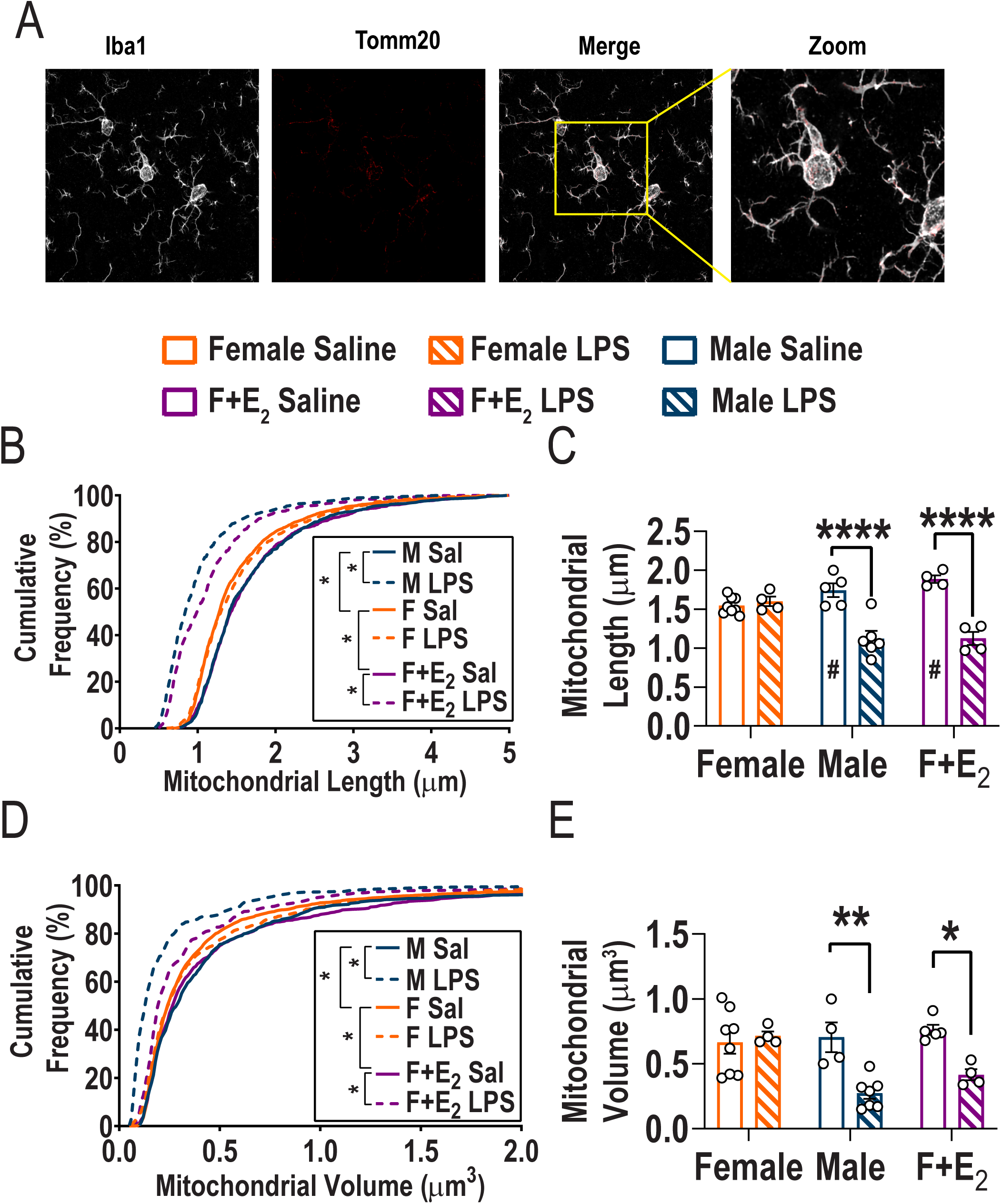
Sex differences in microglial mitochondrial morphometric response to perinatal immune challenge. **A**. Representative images of mitochondria (red) within microglia (grey) in the anterior cingulate cortex of PN30 mice. **B**. Cumulative frequency plots of microglial mitochondrial length analyzed in the ACC of female (orange), male (blue), and masculinized female (F+E_2_: purple) mice. Differences across groups were assessed by Kruskal-Wallis test followed by Dunn’s post-hoc analyses. * p < 0.05. **C**. Average mitochondrial length within microglia in the ACC of female (orange), male (blue), and masculinized female (F+E_2_: purple) mice. Differences across groups were assessed by 2-way ANOVA followed by Bonferonni’s post-hoc analyses. **** p < 0.0001 for male LPS compared to male saline. # p < 0.05 compared to female saline. Bar graphs display mean ± standard error of the mean. **D**. Cumulative frequency plots of microglial mitochondrial volume analyzed in the ACC of female (orange), male (blue), and masculinized female (F+E_2_: purple) mice. Differences across groups were assessed by Kruskal-Wallis test followed by Dunn’s post-hoc analyses. * p < 0.05. **E**. Average mitochondrial volume within microglia in the ACC of female (orange), male (blue), and masculinized female (F+E_2_: purple) mice. Differences across groups were assessed by 2-way ANOVA followed by Bonferonni’s post-hoc analyses. * p < 0.05 ** p < 0.01 for male LPS compared to male saline. # p < 0.05 compared to female saline. Bar graphs display mean ± standard error of the mean.

Mitochondria form dynamic, interconnected networks that are vital to the energetic function of the cell (Bach et al., 2003; Lackner, 2014). In addition to morphometric analyses of individual microglial mitochondrial morphologies, we also assessed the ability of early-life immune challenge to affect connectivity of microglial mitochondrial networks. Mitochondrial connectivity analyses led to similar observations as were seen in morphometric analyses, as PN9 LPS induced a decrease in the number of branches/cell and number of networks/cell in both males and masculinized females, while having no effect on these connectivity measures in female mice (Supplemental Fig. 5C-E). No mitochondrial effects were observed in non-microglia (Iba1 negative cells; Supplemental Fig. 5F-G).

Mitochondrial morphologies are often closely aligned with cellular bioenergetic function (Liesa and Shirihai, 2013; Liesa et al., 2009; Nair et al., 2019; Wai and Langer, 2016). To more closely analyze microglial mitochondrial energetics, we first isolated microglia from the prefrontal cortex of PN10 mice treated 24 hr before with saline or LPS and assessed their oxygen consumption and glucose uptake capacities (Fig. 6A). We assessed cellular oxygen consumption of isolated microglia at baseline as well as following treatment with the protonophore carbonyl cyanide 4-(trifluoromethoxy)phenylhydrazone (FCCP) to stimulate maximal oxygen consumption. There were no significant differences observed in baseline (untreated) oxygen consumption rates (OCR) between microglia from saline or LPS-treated mice, regardless of sex (Fig. 6B-C). In all females as well as saline-treated males, the FCCP-induced maximal OCR was significantly elevated from basal OCR (Fig. 6C). However, FCCP failed to stimulate a significant upregulation in LPS-treated male microglia (Fig. 6C). Importantly, FCCP-stimulated maximal OCR was significantly lower in LPS-treated males compared to saline-treated males and all female groups (Fig. 6C). Similarly, we observed a significant LPS-induced stimulation in glucose uptake from male microglia compared to saline-treated microglia (Fig. 6D). To visualize microglial oxidative and glycolytic energetics, we plotted maximal OCR against glucose uptake, and noted that PN9 LPS treatment does not substantially alter the metabolic phenotype of PFC microglia isolated from females, whereas LPS shifted male microglia towards a more glycolytic phenotype (Fig. 6E).

**Figure 6.**
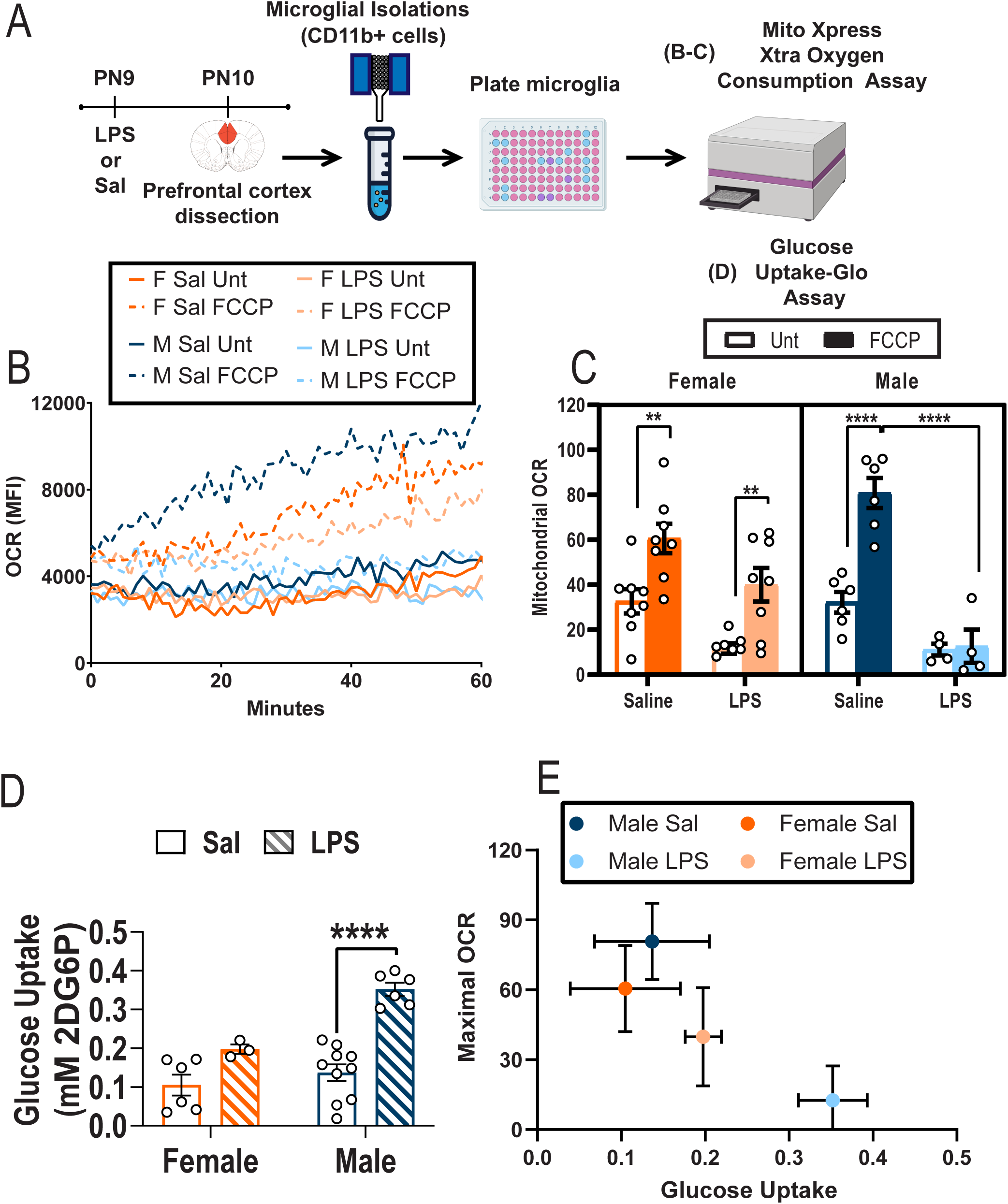
Perinatal immune challenge leads to lasting alterations in mitochondrial function in microglia from male - but not female-mice. **A**. Experimental timeline. Female or male mice were injected on PN9 with LPS or saline vehicle. At PN10, anterior cingulate cortex was dissected, and microglia were isolated by CD11b bead method. CD11b+ cells (microglia) were plated in 96 well plates and cellular oxygen consumption was assessed using Mito Xpress Xtra Oxygen Consumption Assay. Cells were either untreated or maximal oxygen consumption was stimulated by treatment with FCCP prior to sealing of the well. Extracellular acidification rate (ECAR) was assessed using the pH-Xtra Glycolysis Assay. Glucose uptake was assessed using Glucose Uptake-Glo assay. **B**. Representative trace of cellular oxygen consumption rate (OCR) across the 160 minutes of assay run time. Untreated (Unt) cells are depicted by solid lines, whereas FCCP-treated cells are depicted by dashed lines. **C**. Baseline and maximal oxygen consumption rates were assessed. Differences across groups were assessed by 3-way ANOVA followed by Bonferonni’s post-hoc analyses. * p < 0.05. Bar graphs display mean ± standard error of the mean. **D**. Glucose Uptake was assessed using the Glucose Uptake-Glo Assay and reported as mM of 2DG6P uptake. Differences across groups were assessed by 2-way ANOVA followed by Bonferonni’s post-hoc analyses. **** p < 0.0001. Bar graphs display mean ± standard error of the mean. **E**. Maximal oxygen consumption was plotted against Glucose uptake to depict cellular bioenergetic state. Data are presented as mean +/- Standard deviation.

## DISCUSSION

Here we demonstrate the impact of sex on social behavior and microglial mitochondrial responses to early-life immune activation and point to the critical role of gonadal hormones during perinatal development in these sex-biased alterations. Specifically, we show that early life immune activation induces alterations in social behaviors in male but not female mice, and that these behavioral changes are dependent upon microglial Tlr4 signaling. We also demonstrate that early life immune activation reveals male-biased vulnerabilities in mitochondrial gene expression, mitochondrial morphologies, mitochondrial network connectivity, and bioenergetic functions, specifically within microglia. These sex-biased alterations in microglial mitochondrial functions may explain the sex-biased vulnerabilities of social behaviors in response to perinatal immune challenge. Importantly, we induced these behavioral and mitochondrial vulnerabilities to early-life immune activation in chromosomally female mice by masculinizing female pups at birth with estradiol, the aromatized form of gonadally-derived testosterone that can masculinize female brain and behavior (Lenz et al., 2013; McCarthy, 2009; McCarthy et al., 2018). Collectively, these findings provide new evidence that sex-specific perinatal brain development is heavily influenced by gonadal hormones, and that this hormone-dependent development organizes male-biased susceptibility to inflammatory perturbations in the perinatal environment.

We initially hypothesized that early-life immune activation induces male-biased deficits in social behavior through male-biased recruitment of peripheral inflammatory monocytes (CD11b^+^CD45^hi^ cells) into the brain. Indeed, numerous studies have demonstrated that perturbations such as LPS injection or stress induce infiltration of peripheral immune cells into the brain, where these inflammatory cells act to induce behavioral changes (Weber et al., 2017; Wohleb et al., 2013, 2014a, 2014b). However, these studies often fail to examine whether sex differences in infiltration may occur. In contrast to our hypothesis, we demonstrated no sex difference in the infiltration of peripherally derived monocytes into the brain, as PN9 LPS injection induced a robust increase in infiltration into both male and female brains. The lack of observed sex differences in peripheral immune infiltration next led us to assess whether immune signaling within microglia was necessary for the sex-biased behavioral alterations. To do so, we used mice in which Tlr4, a pattern recognition receptor to which LPS binds, was ablated within microglia (Ceasrine et al., 2022). Importantly, ablation of Tlr4 signaling within microglia did not impact sociability or social novelty preference in the absence of an immune challenge. Knocking out Tlr4 signaling within Cx3cr1+ cells prevented LPS-induced alterations, demonstrating the necessity of direct microglial immune signaling in this sex-biased behavioral response. The Cre-LoxP transgenic technology used in this study ablated Tlr4 in all Cx3cr1-expressing cells, i.e. tissue resident macrophages. That the infiltrating monocytes observed following early life immune activation do not express Cx3cr1 further suggests that infiltrating monocytes were not contributing to the male-biased behavioral vulnerability. However, future studies using more precisely targeted genetic ablation strategies will be important to further discern the impact of peripheral vs central macrophage immune signaling in the regulation of social behaviors.

Activation of Tlr4 by LPS results in robust changes in the morphology and function of mitochondria within microglia, and microglial Tlr4-mediated alterations in neuronal functions require alterations in microglial mitochondrial metabolism (Nair et al., 2019; York et al., 2020). We therefore next assessed whether early-life immune activation could be leading to the male bias in social behavioral deficits through sex-biased alterations in microglial mitochondria. Consistent with a growing body of literature demonstrating critical links between microglial inflammatory processes and mitochondrial functions (Bernier et al., 2020a; Nair et al., 2019; York et al., 2020), we observed robust male-specific alterations in microglial mitochondria. For instance, we demonstrated that LPS treatment induced a strong downregulation of mitochondrial ETC genes specifically in microglia from male mice. LPS treatment did not similarly downregulate ETC genes in female microglia, although masculinizing female mice at birth with male-typical gonadal hormones conferred a similar susceptibility to immune activation in ETC genes. This programming of vulnerability appears to be cell-type specific, as CD11b^-^ cells did not show alterations in mitochondrial ETC gene expression in response to LPS treatment. In addition to gene expression, we demonstrate robust sex- and microglia-specific alterations in mitochondrial morphology that are induced by PN9 LPS challenge. Importantly, we demonstrate that the developmental gonadal hormone surge that is necessary for typical male masculinization of the body and brain also results in male microglial mitochondrial morphological vulnerability to early-life immune activation. Together, these results suggest that mitochondria specifically within microglia are particularly vulnerable to immune activation in male mice, and that this vulnerability is organized by the perinatal surge in gonadal hormones that is necessary to typically masculinize the male brain and body.

Our findings identify microglia and mitochondria as potentially important therapeutic targets in the treatment of social alterations in neurodevelopmental disorders. This aligns with previous work in other mouse models as well as postmortem analyses of brains obtained from patients diagnosed with neurodevelopmental disorders such as ASD and SZ that demonstrate altered microglial morphologies and mitochondrial alterations (Morgan et al., 2010; Rosenfeld et al., 2011; Tang et al., 2013; Tetreault et al., 2012). Of particular interest is our finding that the sex-biased downregulation of mitochondrial ETC genes induced by LPS was somewhat specific for genes encoding Complex I of the mitochondrial ETC. Complex I is especially vulnerable to impairment by oxidative stress and is demonstrated to be dysfunctional in a wide variety of diseases. Due to the relative susceptibility of Complex I, particular attention has been paid to the development or repurposing of treatments that target Complex I functions. Our findings raise the possibility that more targeted mitochondrial drugs that can bypass dysfunction in Complex I, such as the electron donor idebenone (Jaber and Polster, 2015), may be particularly useful for the treatment of neurodevelopmental disorders. The recent success of mitochondrial transplantation to restore mitochondrial function across disease and injury models may prove vital to the treatment of sex-biased neurodevelopmental disorders with mitochondrial dysfunction. However, brain region- and cell type-specific targeting of mitochondrial function, through either direct mitochondrial transplantation or other therapeutic regimens directly in the brain, will surely remain difficult in human patients for the foreseeable future. Other indirect avenues to impact brain mitochondrial function, particularly within microglia, will likely prove vital to treatment and prevention of behavioral symptoms associated with neurodevelopmental disorders such as ASD.

In summary, perinatal hormone exposure has been associated with increased risk of offspring neurodevelopmental disorder diagnosis (Baron-Cohen et al., 2020; Bilder et al., 2019), but the mechanisms through which hormone exposure may program male behavioral vulnerability, or provide protection in females, are scarce. We demonstrate that the male-specific perinatal hormone surge that is essential for masculinizing the male brain introduces vulnerability to social behavior deficits in response to early-life immune activation via microglial Tlr4 signaling. Within microglia, we demonstrate that the mitochondrial functional vulnerability to immune challenge, consistent with previous findings, is also particularly vulnerable and highly specific to male mice. This male-biased behavioral and microglial mitochondrial vulnerability is programmed by the male-typical perinatal gonadal hormone surge.

## Limitations of the study

We report that PN9 LPS challenge induced a robust increase in the infiltration of peripheral immune cells into the brains of both male and female mice. The lack of sex difference in infiltrating cell number does not rule out the possibility that the infiltrating cells may act differently between males and females. For instance, infiltrating cells may induce behavioral deficits in male brains whereas infiltrating immune cells could be providing a female protective factor against immune challenge. Additionally, we did not assess whether there were sex differences in the infiltration of other immune cells such as T cells, which have previously been shown to impact microglial function as well as mitochondrial function following their migration into the brain (Fan et al., 2019). Therefore, a deeper understanding of immune dysregulation in the brain, both in terms of type of infiltrating cell as well as functionality of these cells following perinatal perturbations is needed in both females and males.

Microglia play an important role in the development and regulation of neural circuits (Dziabis and Bilbo, 2022), and our lab has recently shown that phagocytosis of dopamine D1 receptors regulate the development of social play behaviors in male but not female rodents (Kopec et al., 2018). Interestingly, recent studies using a mouse model in which the mitochondrial protein UCP2 was ablated specifically within microglia demonstrated male-specific alterations in synaptic architecture and neural functions (Yasumoto et al., 2021), suggesting a link between microglial mitochondrial function and synaptic regulation. These data build upon other findings that mitochondrial dysfunction and altered metabolism play a critical role in the regulation of social behaviors (Kanellopoulos et al., 2020; Miranda Mendonça et al., 2019; Picard et al., 2018), suggesting that altered microglial mitochondrial function may regulate the establishment of complex social behaviors through alterations in synaptic regulation. Previous studies have demonstrated that microglial phagocytosis regulates the development of synaptic circuitry important for social behavior in the ACC during the time period surrounding our PN9 injection (Block et al., 2022). Given the importance of the ACC to regulation of social behaviors, and it’s implication in the etiology of various neurodevelopmental disorders (Block et al., 2022; Delmonte et al., 2013; Mague et al., 2020), we assessed microglial mitochondrial function within the ACC. While our studies revealed robust alterations in microglial mitochondrial function in the ACC, this study did not comprehensively characterize mitochondrial function from every region that may control these complex behaviors. Future studies characterizing immunometabolic responses to inflammatory challenges, and whether these microglial mitochondrial alterations influence synaptic pruning/phagocytosis by microglia, across all brain regions, will be highly informative.

## ACKNOWLEDGEMENTS

This work was supported by R01 ES025549 to SDB, F32 MH116604 to EAB, P50 MH115874 to WAC, Paul and Janis Cunningham, and by the Robert and Donna Landreth Family Foundation.

