## Supplementary Materials for "Gonadal Hormones Impart Male-Biased Behavioral Vulnerabilities to Immune Activation via Microglial Mitochondrial Function"

Staci Bilbo

Duke University, Durham, NC 27708

**This PDF file includes:**

Materials and Methods

Supplementary Text

Figs. S1-S5

### METHODS

#### EXPERIMENTAL MODEL AND SUBJECT DETAILS

##### Experimental Animals

Tlr4-floxed mice were purchased from Jackson Laboratories (Bar Harbor, ME: Stock # 024872). Cx3cr1-CreBT (MW126GSat) mice were generated and provided by L. Kus (GENSAT BAC Transgenic Project, Rockefeller University, NY) and backcrossed over 12 generations on a C57BL/6J (Charles River Laboratories) background. Wild type (WT) C57BL/6J mice were purchased from Jackson Laboratories (Bar Harbor, ME: Stock # 000664). Cx3cr1-CreBT mice were bred with Tlr4-floxed mice over 3 generations to generate offspring that were fully Tlr4-floxed (F/F) and either Cre positive (Tlr4 cKO: removal of microglial Tlr4) or Cre negative (Ctrl: no modification to microglial Tlr4). Transgenic mouse genotyping was performed using polymerase chain reaction on tail-spin DNA. The following primers were used for *Tlr4* genotyping: Forward: TGA CCA CCC ATA TTG CCT ATA C and Reverse: TGA TGG TGT GAG CAG GAG AG, per supplier instructions. All mice were group housed in standard mouse cages under standard conditions (12 hour light/dark cycle, 23°C, 50% humidity) with same-sex littermates. All experiments were performed in accordance with the NIH *Guide to the Care and Use of Laboratory Animals* and approved by the Massachusetts General Hospital Institutional Animal Care and Use Committee.

#### METHOD DETAILS

##### Masculinization

100 ug estradiol benzoate (Sigma-Aldrich catalog # E8515, St. Louis, MO, USA) dissolved in sesame seed oil was administered subcutaneously on postnatal days 1 and 2. This dose and timing were chosen based on previous work (Bowers et al., 2010; VanRyzin et al., 2019b) demonstrating ability to masculinize male rodent body and brain.

**PN9 Saline or lipopolysaccharide Injection**

10 mg/kg lipopolysaccharide (from Escherichia coli serotype 0111:B4, Sigma-Aldrich, St. Louis, MO, USA) dissolved in 0.9% sterile saline or 0.9% sterile saline control was administered subcutaneously on postnatal day 9 (PN9). This dose and timing were chosen based on previous work demonstrating its ability to induce male-specific changes in social behavior (Carlezon et al., 2019). All animals within a litter received the same injection to prevent any indirect exposure to the other drug treatments.

**Behavioral Assays**

Behavioral assays were conducted at PN15 and PN30-45 in a separate cohort of animals from those used for molecular analyses. For all assays, testing took place in the first half of the light phase. Mice were moved to the behavioral testing room 1 hr prior to behavioral testing to habituate to the room each day and were additionally habituated to the testing room for 4 hr prior to the first day of testing. All behavior was video recorded and hand-scored using Solomon Coder (https://solomon.andraspeter.com) by a blinded observer.

**Juvenile Social Exploration**

Social exploratory behavior was assessed at PN15, an age in which mouse pups have opened their eyes and are becoming more independent (Bilbo et al., 2018). Extra nesting material was placed into the home cages of age and treatment-matched litters at PN13. On the day of testing, litters are acclimated to testing room for 1 hr prior to testing. A quarter-sized amount of nesting material is taken from the cage of the subject mouse and placed on one side of the testing cage that contains clean bedding material. A quarter-sized amount of nesting material is also taken from the cage of an age- and treatment-matched litter and spread on the opposite side of the testing cage. PN15 mice are then placed in the center of the cage and allowed to explore for 3 min. For scoring purposes, the cage was divided into three areas: home nesting, center, and unfamiliar nesting. The amount of time spent in the home nesting side, and the latency to first enter the side containing unfamiliar litter nesting was quantified.

**Sociability and Social Novelty Preference Tasks**

A 3-chambered social preference task was used to test the preference of mice to investigate either a social stimulus or a non-social stimulus (sociability) or a novel social stimulus vs a familiar social stimulus (social novelty preference), as previously described (Smith et al., 2020). For the sociability task, novel age-, sex- and treatment-matched conspecific mice or a novel rubber duck were placed in small plexiglass containers on opposite sides of the 3-chambered box. Experimental mice were placed in the center chamber and allowed to freely explore for 5 min. Separate 3-chambered boxes were used for female and mice male, and sexes were tested on separate days. Sociability was calculated as the amount of time spent investigating the novel social mouse as a proportion of total time spent investigating any stimulus as a quantification of social preference. For the social novelty preference task, a similar procedure was used, with the two stimuli this time consisting of a new novel age-, sex- and treatment-matched conspecific mouse or a familiar cage mate sibling. Experimental mice were then placed in the center chamber and allowed to freely explore for 10 min. The following behavioral outcomes were manually scored using Solomon Coder: time spent investigating the novel mouse, time spent investigating the novel object (sociability task), time spent investigating the familiar cage mate sibling (social novelty preference task), and time spent in each chamber.

**Elevated Zero Maze**

The elevated zero maze was used to test for anxiety-like behavior as previously described (Bolton et al., 2017). We used a zero maze (Stoelting) with an elevated (50 cm height) circular lane (5 cm wide) that is divided into four quadrants. Two quadrants are enclosed by 15 cm tall walls with the remaining two quadrants left exposed. Test mice were placed into a closed arm, and time spent in the open or closed quadrant arms was scored over a 5 min period.

**Light Dark Box**

The light dark box task was used as a second test for anxiety-like behavior as previously described. We used a box with 35 cm tall walls that was split into two 20cm x 40cm chambers. The walls on one side were clear and brightly illuminated while the walls of the other chamber were opaque and the chamber itself was dark, with a small open space allowing for free passage between chambers. Test mice were placed into the dark chamber, and time spent in the light or dark chambers was scored over a 10 min period.

**Marble Burying**

Standard mouse cages were prepared with 5 cm depth wood shavings (5 cm depth). 20 black and blue colored marbles were arranged on the top of the wood shaving in a 5x4 grid formation. Mice were then placed in the cage for 20 min, after which mice were removed from cage and a picture was taken. A blinded observer assessed number of buried marbles. Marbles were considered to be buried if > 2/3 of surface was no longer visible above the wood shavings.

**Microglia isolation for gene expression**

Microglia isolations were performed as previously described (Bordt et al., 2020b). CD11b+ cells (herein referred to as microglia) were isolated from prefrontal cortex (PFC) on PN30. Following transcardial perfusion with ice-cold saline, PFC was dissected from whole brain on ice using sterile forceps and minced on a petri dish set on ice using a sterile razor blade. Tissue homogenate was then placed in Hank's Buffered Salt Solution (HBSS; ThermoFisher Scientific, NY, USA) with collagenase A (Roche, Indianapolis, IN, USA; Catalog # 11088793001) and 0.4 mg/mL DNase I (Roche, Indianapolis, IN, USA; Catalog # ) and incubated for 15 min at 37°C. Mechanical dissociation of tissue homogenate was performed by pipetting samples through successively smaller flame-polished Pasteur pipettes until a single-cell suspension was obtained. Samples were then filtered, rinsed with HBSS, and centrifuged. Samples were then incubated on ice for 15 min with CD11b-conjugated microbeads (Miltenyi Biotec, San Diego, CA, USA: Catalog # 130-093-634), and passed through a magnetic bead column (Miltenyi Biotec, San Diego, CA, USA: Catalog # 130-042-401) to separate CD11b+ and CD11b- cells. Cells were washed in 1X PBS and resuspended in Trizol (ThermoFisher Scientific, NY, USA: Catalog # 15596026), after which they were stored at -80°C until RNA extraction.

**RNA extraction**

Isolated microglia were homogenized in Trizol, mixed for 10 min at 2000 rpm on a MixMate (Eppendorf), followed by resting at room temperature for 15 min. Chloroform (1:5 with Trizol) was then added and samples were mixed for 2 min at 2000 rpm on a MixMate (Eppendorf). Tubes then sat at room temperature for 3 min followed by centrifugation for 15 min at 11,900 rpm at 4°C. Aqueous phase was aliquoted into a fresh tube and isopropanol was added (1:1 with aqueous phase volume) along with 2 uL of GlycoBlue coprecipitant (ThermoFisher Scientific: Catalog # AM9515). Tubes were then mixed for 1 min at 2000 rpm on a MixMate, allowed to incubate at RT for 10 min, and centrifuged at 11,900 rpm for 10 min at 4°C. Pellets were then rinsed twice in ice-cold ethanol (75%) and resuspended in 6-8 uL of nuclease-free water (ThermoFisher Scientific: Catalog # AM9932). RNA quantity and purity was assessed using a NanoDrop 2000 (ThermoFisher Scientific).

**cDNA synthesis and qPCR**

cDNA was synthesized from 200 ng of total RNA using the QuantiTect Reverse Transcription Kit (Qiagen, Hilden, Germany) according to manufacturer's instructions. qPCR was subsequently performed using this cDNA and Taqman gene expression assays performed on a QuantStudio 5 Real-Time PCR System (ThermoFisher Scientific) as previously reported (Bordt et al., 2021). Gene expression was normalized to the reference gene 18S and calculating using 2-ΔΔCt method relative to the lowest sample on the plate.

**cDNA synthesis and PCR Array**

cDNA to be used for PCR Array was synthesized from 200 ng of total RNA using the $RT^2$ First Strand Kit (Qiagen: Catalog # 330404) according to manufacturer's instructions. Mitochondrial gene expression was then assessed using the $RT^2$ Profiler™ PCR Array Mouse Mitochondrial Energy Metabolism (Qiagen: Catalog # PAMM-008ZA-24) and $RT^2$ SYBR Green ROX qPCR Mastermix (Qiagen: Catalog # 330523) according to manufacturer's instructions.

**Immunohistochemistry and Confocal Microscopy**

Following transcardial perfusion with ice-cold saline followed by 4% paraformaldehyde (PFA), brains were post-fixed in 4% PFA for 48 hr at 4°C. PFA-fixed brains were then cryoprotected in 30% sucrose for at least 48 hr at 4°C. Brains were then flash frozen in 2-methylbutane on dry ice and stored at -80°C before cryosectioning at 30 μm on a Leica cryostat. Brain sections were stored in cryoprotectant until staining. Tissue sections of interest were then removed from cryoprotectant and rinsed 3x 5 min in 1X PBS to remove excess cryoprotectant. Tissue was first permeabilized in 0.3% Triton-X100 in PBS for 30 min and rinsed 3x 5 min in 1X PBS. Epitope retrieval was then performed by incubating at 80°C in 10 mM citric acid (pH 9.0) for 30 min followed by 3x 10 min 1X PBS washes. Background fluorescence was then quenched by 60 min incubation in 1 mg/mL sodium tetraborate in 0.1M PB, followed by 6x 5 min PBS washes and incubation in 50% methanol for 60 min followed by 6x 5 min PBS washing. Sections were then blocked in 10% normal goat serum (Vector Laboratories: Catalog # S-100-20) in PBS for 60 min. Primary antibodies were applied sequentially for overnight incubations in 5% normal goat serum and 0.3% Tween20 in PBS at 4°C. Primary antibodies used were chicken anti-Iba1 (Synaptic Systems 234 006, 1:500) and rabbit anti-Tomm20 (Novus NBP2-67501, 1:500). Secondary antibodies were then applied for 2 hr at RT protected from light in 5% NGS with 0.3% Tween20 followed by 6x 10 min PBS washes. Secondary antibodies used were goat anti-rabbit highly cross-adsorbed Alexa Fluor Plus 488 (ThermoFisher Scientific A32731, 1:200) and goat anti-chicken Alexa Fluor 568 (ThermoFisher Scientific A-11041, 1:200). Sections were then mounted on gelatin subbed slides, coverslipped with VECTASHIELD Antifade Mounting Medium with DAPI (Vector Laboratories, Burlingame, CA, USA: Catalog #H-1000), sealed with nail polish, and protected from light until imaging.

**Imaging Analyses**

To quantify immunofluorescence of Iba1 and Tomm20 staining, 63x (x1.39) magnification Z-stacks were taken on a Nikon A1R confocal microscope. Z-stacks were taken every 0.2 μm. Mitochondrial length and volume within microglia was quantified using Imaris software (Bitplane). Individual channel backgrounds (microglia, mitochondria, nuclei) were first subtracted and then threshold values were recorded using Fiji software. Images were then uploaded into Imaris to create volumetric images using the Surface tool. Mitochondria were masked (Kopec et al., 2018) inside of the microglial (Iba1+) signal prior to reconstruction in order to quantify mitochondrial morphology within microglia, and then mitochondrial length was measured using the BoundingBoxOO Length C setting (longest principal axis) as previously described (Chandra et al., 2017). All Tomm20 signal masked outside of microglia (Iba1-) was used to quantify non-microglial signal. Mitochondrial volume was also exported for statistical analysis.

**Flow Cytometry**

Mice were transcardially perfused with ice-cold saline until perfusate was clear, after which prefrontal cortex was dissected on ice and minced on a petri dish set using a sterile razor blade. Tissue homogenate was then placed in Hank's Buffered Salt Solution (HBSS; ThermoFisher Scientific, NY, USA) with collagenase A (Roche, Indianapolis, IN, USA; Catalog # 11088793001) and 0.4 mg/mL DNase I (Roche, Indianapolis, IN, USA; Catalog # ) and incubated for 15 min at 37°C. Mechanical dissociation of tissue homogenate was performed by pipetting samples through successively smaller flame-polished Pasteur pipettes until a single-cell suspension was obtained. Samples were then filtered, rinsed with HBSS, and centrifuged. Cells were then cleaned using Debris Removal Solution (Miltenyi Biotec: Catalog # 130-109-398), washed with PBS, and then plated in a 96 well U-bottomed plate at 70,000 cells/well in Stain Buffer (BD Biosciences: Catalog # 554657). Cells were blocked with TruStain FcX (anti-CD16/32) antibody (1:100; BioLegend: Catalog # 101320) and stained with LIVE/DEAD Fixable Blue Dead Cell Stain Kit (ThermoFisher Scientific: Catalog # L23105). Cells were then labelled with anti-CD11b-BV510 (1:100; BD Biosciences: Catalog # 562950), anti-CD45-APC/Cy7(1:200; BioLegend: Catalog # 103116), anti-CCR2-APC (1:100; R&D Systems: Catalog # FAB5538A-100), and anti-Cx3cr1-FITC (1:100; BioLegend: Catalog # 149019) antibodies in Stain Buffer. Each experiment contained a fluorescence minus one (FMO) control and was analyzed on a Beckman Coulter CytoFLEX flow cytometer. Compensation was performed using VersaComp Antibody Capture Kit (Beckman Coulter: Catalog # B22804) according to manufacturer's instructions.

In this study, we chose to isolate microglial cells from the brain using a magnetic bead enrichment method (Bordt et al., 2020b) over isolation by fluorescence activated cell sorting (FACS) due to the known ability of FACS to upregulate immediate early gene expression (Ayata et al., 2018; Haimon et al., 2018; Li et al., 2019), induce transcription factors associated with proinflammatory states (Ayata et al., 2018), and to alter cellular redox state (Llufrio et al., 2018). However, an obvious caveat of our CD11b-based enrichment strategy is the single-antigen approach, which is likely to pull down immune cells present in the brain, not only microglia. Our findings that there were not sex differences in the number of $CD11b^{+}CD45^{hi}$ cells infiltrating into the brain following suggest that any sex difference found in isolated cell biology would be due to microglia, but this is a limitation of our study. Future studies of microglial biology could be performed using FACS and newly-published transcriptional inhibitors to dampen some of the negative impacts of cell sorting (Marsh et al., 2022).

#### QUANTIFICATION AND STATISTICAL ANALYSES

##### Statistical Analysis

All statistics and data visualization were performed using GraphPad Prism Version 9. Exact statistical tests performed for each experiment are reported in the figure legends. Individual points on graphs represent individual biological samples. Statistical significance was determined at $p < 0.05$.

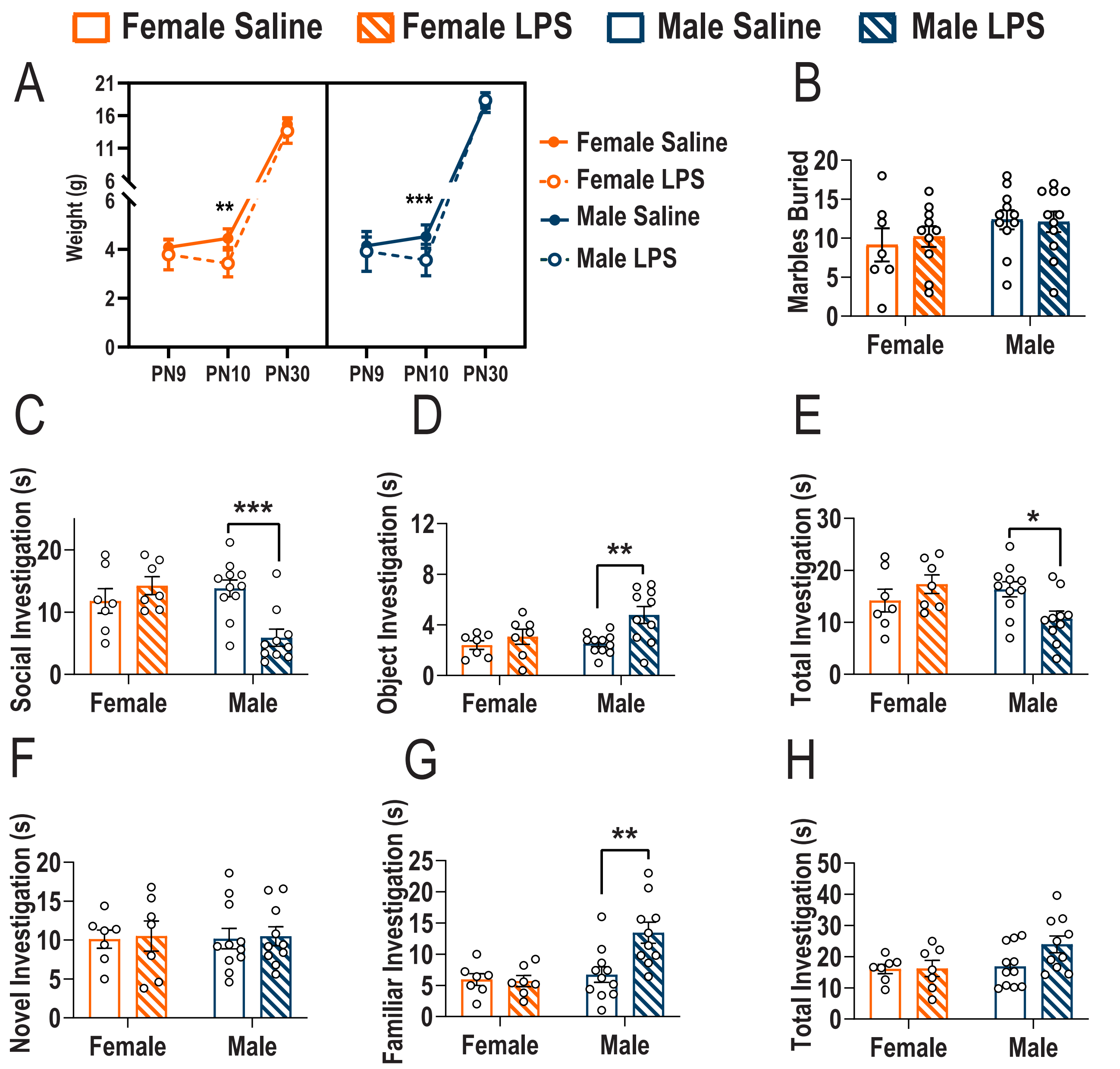


**Supplemental Figure 1. Extended sex-biased behavioral responses.**
**A.** Female (orange) and male (blue) mice offspring challenged with saline (solid lines, closed circles) or LPS (striped bars, open circles) at PN9 were weighed at PN9 prior to injection, at PN10 (24 hr post injection), and at PN30 (prior to behavioral testing). Differences across groups were assessed by 3-way ANOVA followed by Bonerroni's post-hoc analyses. Significant differences were observed only at PN10. ** $p < 0.01$. *** $p < 0.0001$.
**B.** PN30 female (orange) and male (blue) mice challenged with saline (Sal: open bars) or LPS (striped bars) at PN9 were placed in a cage with 20 marbles for 20 min, after which number of marbles buried was assessed. No significant differences across groups were found by 2-way ANOVA.
**C-E.** PN30 female (orange) and male (blue) mice challenged with saline (Sal: open bars) or LPS (striped bars) at PN9 were placed in a 3 chambered arena and given the choice to interact with a novel conspecific mouse or an inanimate object for 5 min. Time spent investigating the (C) novel animal, (D) novel object, and (E) total investigation time was assessed. Note that raw investigation values are reported in Fig. 1C and are reported in this Supplement to enable a different statistical comparison. Differences across groups were assessed by 2-way ANOVA followed by Bonferonni's post-hoc analyses. * $p < 0.05$ ** $p < 0.01$ *** $p < 0.001$.
**F-H.** PN30 female (orange) and male (blue) mice challenged with saline (Sal: open bars) or LPS (striped bars) at PN9 were placed in a 3 chambered arena and given the choice to interact with a novel conspecific mouse or a cage mate for 10 min. Time spent investigating the (F) novel animal, (G) familiar animal, and (H) total investigation time was assessed. Note that raw investigation values are reported in Fig. 1F and are reported in this Supplement to enable a different statistical comparison.Differences across groups were assessed by 2-way ANOVA followed by Bonferonni's post-hoc analyses. ** $p < 0.01$.

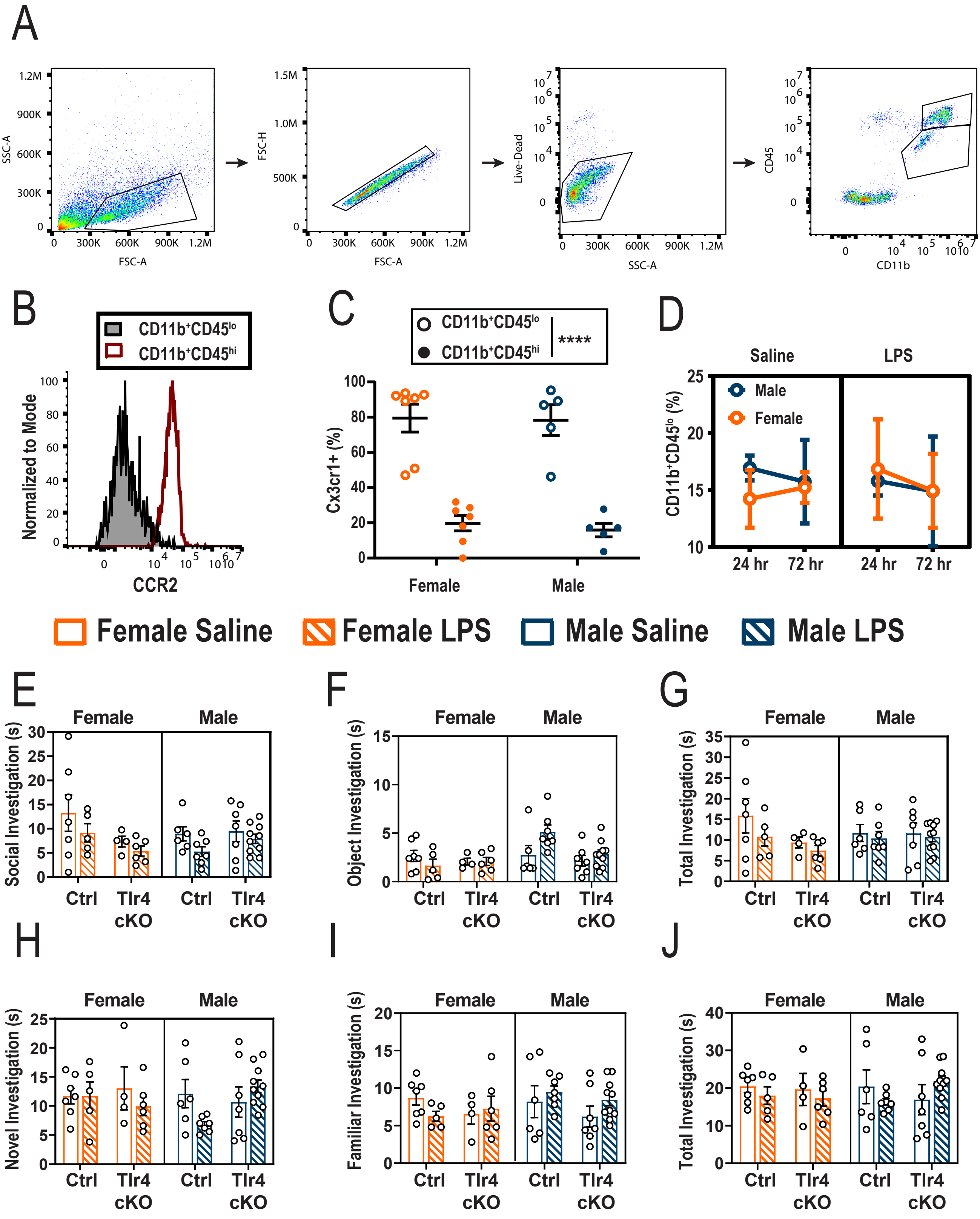


**Supplemental Figure 2. Extended flow cytometry and Tlr4 social behavior results.**
**A.** Prefrontal cortex from female and male mice injected on PN9 with saline or LPS and collected either 24 hr or 72 hr post-infection. Tissue was dissociated, brought to a single cell suspension, and cleamed with Miltenyi Debris Removal Solution prior to flow cytometric analysis. Gating strategy as cells were gated on forward scatter (FSC)/side scatter (SSC), singlets, live-dead, CD11b, and CD45.
**B.** CCR2 expression was assessed with CD11b$^{+}$CD45$^{lo}$ and CD11b$^{+}$CD45$^{hi}$ populations.
**C.** CD11b$^{+}$ cells that were either CD45$^{lo}$ or CD45$^{hi}$ were assessed for Cx3cr1 positivity. Differences across groups were assessed by 2-way ANOVA. A main effect of CD45 hi/lo status (* $p < 0.0001$) was found.
**D.** Brain cells from PN10 and PN12 male and female mice injected on PN9 with saline or LPS were analyzed for CD11b positivity and CD45 hi/lo status. The percentage of cells assessed to be CD11b$^{+}$CD45$^{lo}$ is presented.
**E-G.** PN30 female (orange) and male (blue) Tlr4flox/flox (Ctrl) and Cx3cr1-CrebT:Tlr4flox/flox (Tlr4 cKO) mice challenged with saline (Sal: open bars) or LPS (striped bars) at PN9 were placed in a 3 chambered arena and given the choice to interact with a novel conspecific mouse or an inanimate object for 5 min. Time spent investigating the (E) novel animal, (F) novel object, and (G) total investigation time was assessed. Differences across groups were assessed by 3-way ANOVA followed by Bonferonni's post-hoc analyses.
**H-J.** PN30 female (orange) and male (blue) Tlr4flox/flox (Ctrl) and Cx3cr1-CrebT:Tlr4flox/flox (Tlr4 cKO) mice challenged with saline (Sal: open bars) or LPS (striped bars) at PN9 were placed in a 3 chambered arena and given the choice to interact with a novel conspecific mouse or a cage mate for 10 min. Time spent investigating the (H) novel animal, (I) familiar animal, and (J) total investigation time was assessed. Differences across groups were assessed by 3-way ANOVA followed by Bonferonni's post-hoc analyses.

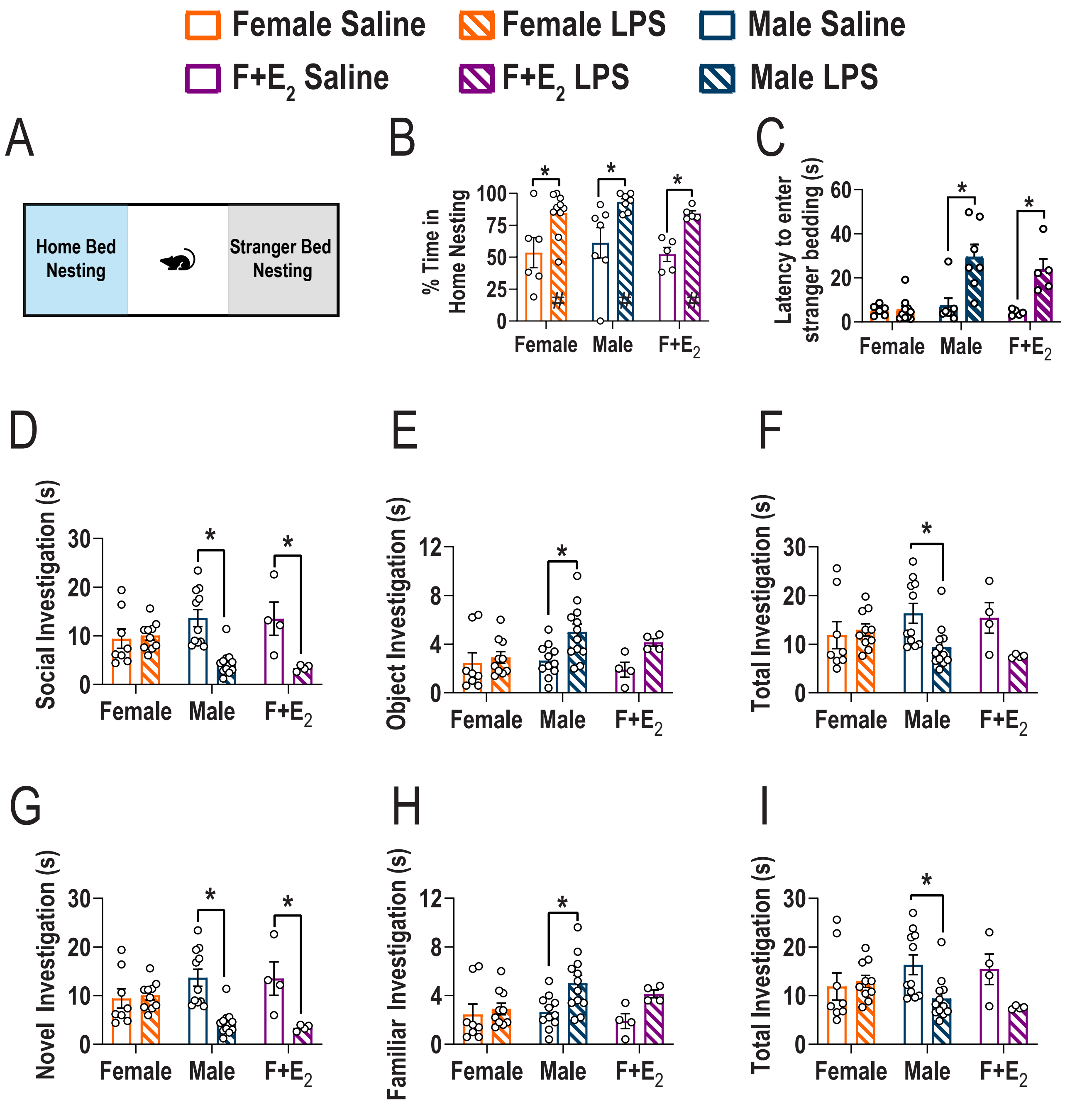


**Supplemental Figure 3. Extended masculinization behavioral analyses.**
**A-C.** PN15 pups were placed into the center of a novel cage with nesting material from their home cage placed on one side and nesting material from an age- and treatment-matched cage placed on the other side. The time spent **(B)** exploring the home nesting side and **(C)** latency to enter the side containing stranger bedding were assessed.
**D-F.** PN30 female (orange) and male (blue) mice challenged with saline (Sal: open bars) or LPS (striped bars) at PN9 were placed in a 3 chambered arena and given the choice to interact with a novel conspecific mouse or an inanimate object for 5 min. Time spent investigating the (D) novel animal, (E) novel object, and (F) total investigation time was assessed. Differences across groups were assessed by 2-way ANOVA followed by Bonferonni's post-hoc analyses. * $p < 0.05$ ** $p < 0.01$ *** $p < 0.001$.
**G-I.** PN30 female (orange) and male (blue) mice challenged with saline (Sal: open bars) or LPS (striped bars) at PN9 were placed in a 3 chambered arena and given the choice to interact with a novel conspecific mouse or a cage mate for 10 min. Time spent investigating the (G) novel animal, (H) familiar animal, and (I) total investigation time was assessed. Differences across groups were assessed by 2-way ANOVA followed by Bonferonni's post-hoc analyses. ** $p < 0.01$.

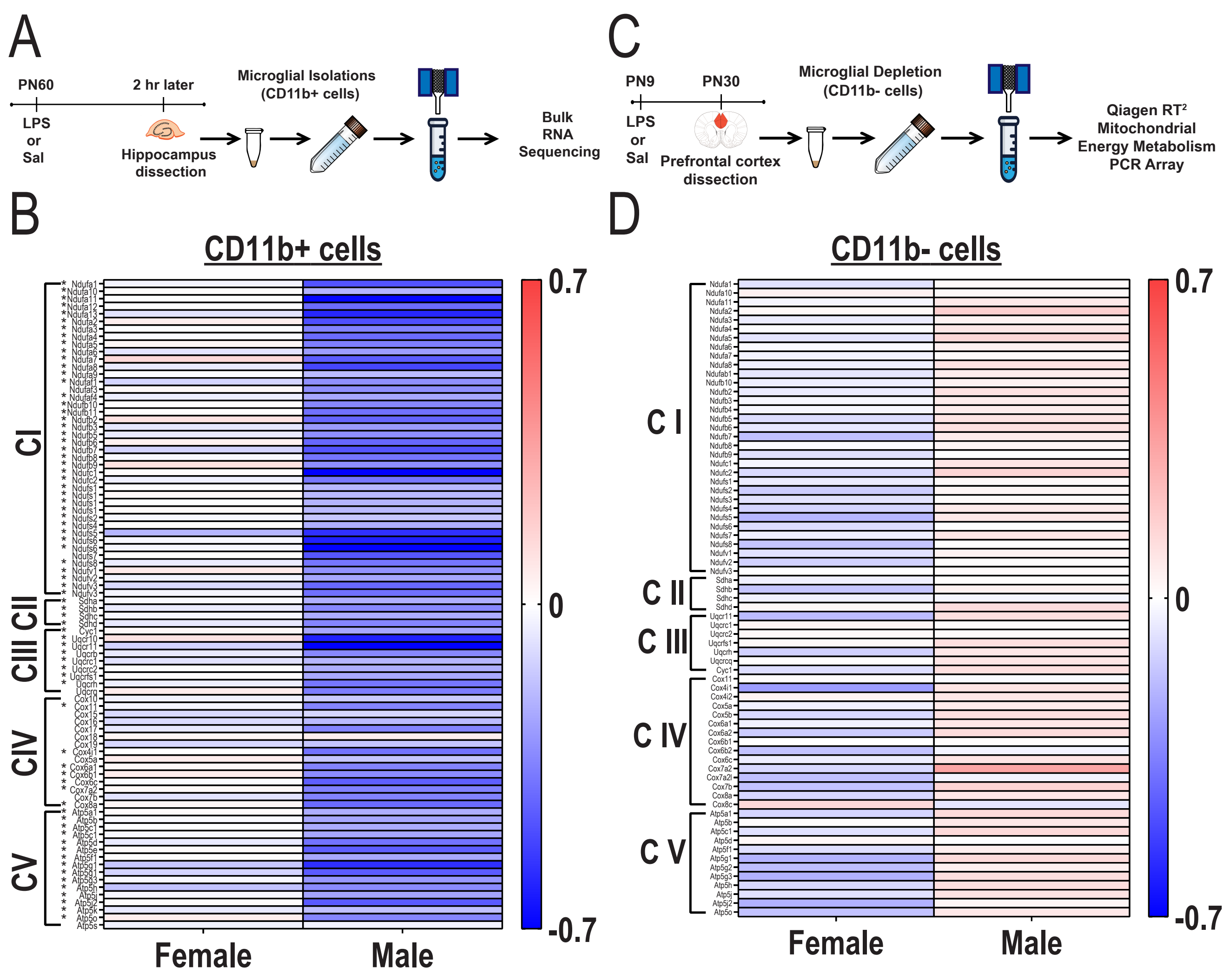


**Supplemental Figure 4. Immune challenge induces acute changes in expression of microglial mitochondrial electron transport chain genes in male - but not in female - mice while not affecting non-microglial cells.**

**A.** Experimental timeline. Female or male mice were injected on PN60 with LPS or saline vehicle. 2 hr later bilateral hippocampus was dissected, and microglia were isolated by CD11b bead method. Extracted RNA from CD11b+ cells (microglia) was then assessed by RNA sequencing, and expression of mitochondrial electron transport chain (ETC) subunits was assessed.

**B.** Log of mRNA expression of ETC subunits (Complexes I-V: CI-CV) from CD11b+ cells isolated from PN60 mice injected 2 hr prior with LPS on PN9 relative to that gene's expression from saline-treated mice (red = upregulation, white = no change, blue = downregulation). Differences across groups were assessed by 2-way ANOVA followed by Bonferonni's post-hoc analyses. * $p < 0.05$ for male LPS compared to male saline. There were no LPS-induced differences in female mice.

**C.** Experimental timeline. Female or male mice were injected on PN9 with LPS or saline vehicle. At PN30, anterior cingulate cortex was dissected, and non-microglial cells were isolated by CD11b negative bead selection. Extracted RNA from CD11b- cells (non-microglial cells) was then assessed for mitochondrial electron transport chain (ETC) gene expression using Qiagen RT$^2$ Mitochondrial Energy Metabolism PCR Array.

**D.** Log of mRNA expression of ETC subunits (Complexes I-V: CI-CV) from CD11b- cells isolated from ACC of PN30 mice injected with LPS on PN9 relative to that gene's expression from saline-treated mice (red = upregulation, white = no change, blue = downregulation). Differences across groups were assessed by 2-way ANOVA followed by Bonferonni's post-hoc analyses.

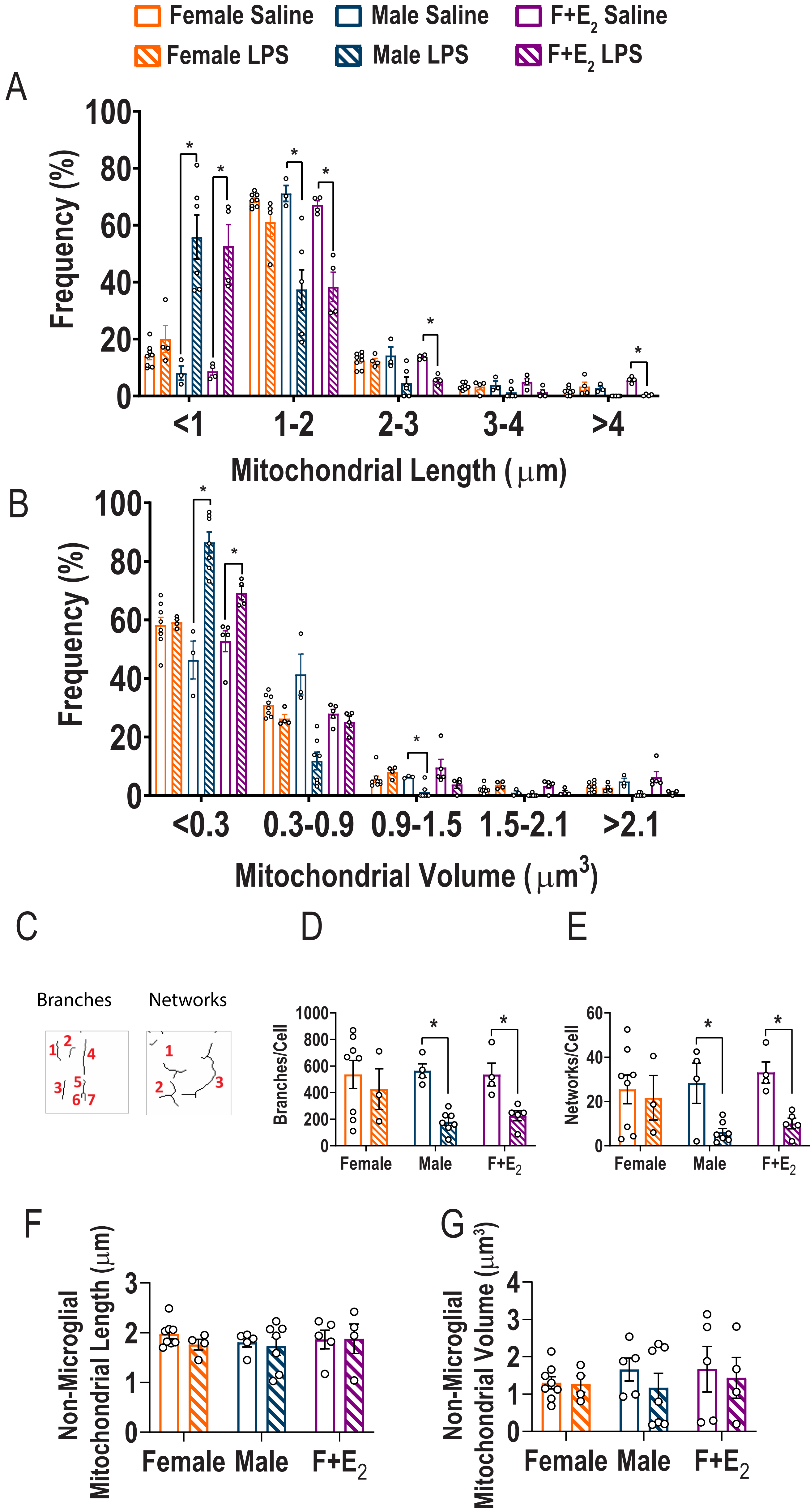


**Supplemental Figure 5. Extended Mitochondrial Morphological Analyses.**
**A-B.** Binned analyses of (A) mitochondrial lengths and (B) mitochondrial volumes. Differences across groups were assessed by mixed model 2-way ANOVA followed by Bonferonni's post-hoc analyses. * $p < 0.05$.
**C.** Skeletons of mitochondrial Tomm20 signal masked inside of microglia (Iba1) generated in Imaris were analyzed for number of branches and number of networks (defined as >2 branches >1 junction).
**D-E.** The number of mitochondrial (D) branches per cell and (E) networks per cell was assessed.
**F.** Average mitochondrial length in cells that are not microglia (masked outside of Iba1) in the ACC of female (orange), male (blue) and F+$E_2$ (purple) mice. No significant differences were observed by 2-way ANOVA.
**G.** Average mitochondrial volume in cells that are not microglia (masked outside of Iba1) in the ACC of female (orange), male (blue) and F+$E_2$ (purple) mice. No significant differences were observed by 2-way ANOVA.
